# Toxoplasma bradyzoites exhibit physiological plasticity of calcium and energy stores controlling motility and egress

**DOI:** 10.1101/2021.05.17.444531

**Authors:** Yong Fu, Kevin M. Brown, Nathaniel G. Jones, Silvia N. J. Moreno, L. David Sibley

## Abstract

*Toxoplasma gondii* has evolved different developmental stages for disseminating during acute infection (i.e. tachyzoites) and for establishing chronic infection (i.e. bradyzoites). Calcium ion (Ca^2+^) signaling tightly regulates the lytic cycle of tachyzoites by controlling microneme secretion and motility to drive egress and cell invasion. However, the roles of Ca^2+^ signaling pathways in bradyzoites remain largely unexplored. Here we show that Ca^2+^ responses are highly restricted in bradyzoites and that they fail to egress in response to agonists. Development of dual-reporter parasites revealed dampened calcium responses and minimal microneme secretion by bradyzoites induced in vitro or harvested from infected mice and tested ex vivo. Ratiometric Ca^2+^ imaging demonstrated lower Ca^2+^ basal levels, reduced magnitude, and slower Ca^2+^ kinetics in bradyzoites compared with tachyzoites stimulated with agonists. Diminished responses in bradyzoites were associated with down-regulation of calcium ATPases involved in intracellular Ca^2+^ storage in the endoplasmic reticulum (ER) and acidocalcisomes. Once liberated from cysts by trypsin digestion, bradyzoites incubated in glucose plus calcium rapidly restored their intracellular Ca^2+^ and ATP stores leading to enhanced gliding. Collectively, our findings indicate that intracellular bradyzoites exhibit dampened Ca^2+^ signaling and lower energy levels that restrict egress, and yet upon release they rapidly respond to changes in the environment to regain motility.

## Introduction

*Toxoplasma gondii* is an obligate intracellular parasite, capable of infecting nearly all warm-blooded animals and frequently causing human infections [1]. The ingestion of tissue cysts in undercooked meat or shed oocysts by infected cats are the major transmission routes of *T. gondii* [2,3]. Following oral ingestion of bradyzoites within tissue cysts or sporozoites within oocysts, the parasite migrates across the intestinal epithelial barrier and disseminates throughout the body as the actively proliferating tachyzoite form that infects many cell types but primarily traffics in monocytes [4]. In response to immune pressure, the parasite differentiates to asynchronously growing bradyzoites within cysts that can persist as chronic infections in muscle and brain tissues [5–7].

Tachyzoites are adapted for rapid proliferation and dissemination due to an active lytic cycle that is controlled at numerous stages by intracellular calcium ion (Ca^2+^) signaling [8]. Artificially elevating intracellular Ca^2+^ using ionophores triggers secretion of microneme proteins, which are needed for substrate and cell attachment, and hence critical for both gliding motility and cell invasion [9–11]. Increase of cytosolic Ca^2+^ released from internal stores is sufficient to trigger microneme secretion [12], and necessary for host cell invasion [12,13], although these processes are also enhanced by the presence of extracellular Ca^2+^ [14]. Increases in intracellular Ca^2+^ also precede egress and drive secretion of perforin like protein 1 (PLP1) from microneme to facilitate rupture of parasitophorous vacuole membrane (PVM) followed by egress [15]. Calcium signaling is initiated by cyclic guanosine monophosphate (cGMP)-generating guanylate cyclase (GC) [16–18] that activates parasite plasma membrane-associated protein kinase G (PKG) [19], stimulating the production of inositol triphosphate (IP_3_) by phosphoinositide-phospholipase C (PI-PLC) and leading to subsequent release of intracellular Ca^2+^ [12,20,21]. Recent studies in *Plasmodium* also implicate PKG in directly controlling calcium through interaction with a multimembrane spanning protein that may function as a channel that mediates calcium release [22]. In turn, Ca^2+^ activates downstream Ca^2+^ responsive proteins including Ca^2+^ dependent protein kinases such as CDPK1 [8] and CDPK3 [23,24], and C2 domain-containing Ca^2+^ binding proteins [25], and calcium binding orthologues of calmodulin [26], which are required for invasion and egress by tachyzoites. Following invasion, protein kinase A catalytic domain 1 (PKAc1) dampens cytosolic Ca^2+^ by suppressing cGMP signaling and reducing Ca^2+^ uptake [27,28]. Collectively, the lytic life cycle of tachyzoites is orchestrated spatially and temporally by controlling levels of intracellular Ca^2+^ and cyclic nucleotides [29].

*Toxoplasma* has evolved elaborate mechanism to control intracellular Ca^2+^ levels through the concerted action of calcium channels, transporters, and Ca^2+^ pumps expressed at the PM and intracellular stores [8,30]. Orthologues to voltage-dependent Ca^2+^ channels, transient receptor potential (TRP) channels, and plasma membrane type Ca^2+^-ATPases (PMCAs) are predicted to be present in *T. gondii* and likely involved in regulating cytosolic Ca^2+^ influx and efflux [31,32]. The endoplasmic reticulum (ER) is the most important storage site from which Ca^2+^ is released to stimulate motility and egress of *Toxoplasma* [8]. SERCA-type Ca^2+^ ATPase is the known mechanism for Ca^2+^ uptake by the ER and its activity, which is inhibited by thapsigargin [33], leads to accumulation of Ca^2+^ in the ER, which when released activates microneme secretion and motility [34,35]. TgA1 a plasma membrane type Ca^2+^ ATPase, transport Ca^2+^ to the acidocalcisome [36], which likely provides a Ca^2+^ sink albeit one that may not be as readily mobilizable as the ER. In addition to internal Ca^2+^ stores, intracellular and extracellular *T. gondii* tachyzoites are capable of taking up Ca^2+^ from host cells and the extracellular environment, respectively, to enhance Ca^2+^ signaling pathways [14,37]. A variety of fluorescent Ca^2+^ indicators that have been developed to directly image Ca^2+^ signals in live cells include Ca^2+^ responsive dyes and genetically encoded indicators [38]. Indicators like Fluo-4/AM, and related derivatives, have been previously used to monitor Ca^2+^ levels in extracellular parasites [34,39]. Genetically encoded calcium indicators such as GCaMP5, GCaMP6f and GCaMP7 have also been used to visualize dynamic Ca^2+^ signals of both intracellular and extracellular tachyzoites with high resolution and sensitivity [37,40–42].

In contrast to tachyzoites, little is known about the roles of Ca^2+^ signaling in control of microneme secretion, gliding motility, and egress by bradyzoites. Although bradyzoites divide asynchronously, they undergo growth, expansion, and sequential rounds of tissue cyst formation and rupture that maintain chronic infection in vivo [5]. Histological studies in animal models support a model of periodic cyst rupture [43], releasing bradyzoites that reinvade new host cells to generate secondary daughter cysts [44], or transition back to actively replicating tachyzoites [45]. Development of bradyzoites has been studied in vitro using systems that induce development due to stress induced by alkaline pH [46] or in cell lines where development occurs spontaneously [47,48]. Although numerous studies have focused on the determinants that control stage conversion between tachyzoites and bradyzoites [6,49], few studies focus on the signaling pathways that control the bradyzoite lytic cycle.

In the present study, we combined stage-specific bradyzoite fluorescent reporters with Ca^2+^ imaging probes to explore Ca^2+^ signaling, microneme secretion, motility and egress by bradyzoites. Our findings indicate that bradyzoites exhibit dampened Ca^2+^ levels, reduced microneme secretion, and minimal egress in response to Ca^2+^ agonists. Ratiometric Ca^2+^ imaging demonstrated lower Ca^2+^ basal levels and significantly lower stored Ca^2+^ in ER and acidocalcisome in bradyzoites, associated with reduced expression of Ca^2+^ ATPases responsible for maintaining intracellular stores. Incubation of extracellular bradyzoites in Ca^2+^ plus glucose lead to rapid recover of both intracellular Ca^2+^ and ATP levels and restored motility. Collectively our findings support a dampened lytic cycle in bradyzoites, arising from diminished Ca^2+^ signaling and lowered energy stores, and that upon release they exhibit rapid metabolic responsiveness to environmental conditions.

## Results

### Ca^2+^ signaling triggers inefficient egress by bradyzoites

To define egress by bradyzoites, we induced the differentiation of tachyzoites to bradyzoites by culture in HFF cells at alkaline pH (8.2) for 7 days. We treated both tachyzoite cultures and in vitro differentiated cysts with Ca^2+^ ionophore A23187 to trigger egress from parasitophorous vacuoles (PVs) or bradyzoite cysts, as detected by indirect immunofluorescence assay (IFA) or time lapse video microscopy. We observed that A23187 induced complete egress of tachyzoites from disrupted PVs while only few bradyzoites were released from cysts that remained largely intact (**Figure 1A**). This result was also confirmed by time-lapse video microscopy using the ME49 BAG1-mCherry strain either grown as tachyzoites (**Figure 1-video 1**) or bradyzoites (**Figure 1-video 2**). We quantified the percentage of tachyzoites or bradyzoites that were released during egress in response to A23187 or the agonist zaprinast, which is a cGMP specific phosphodiesterase (PDE) inhibitor that activates PKG-mediated Ca^2+^ signaling, leading to egress. In contrast to tachyzoites, we found significantly lower egress rate of bradyzoites in response to A23817 or zaprinast (**Figure 1B**). To examine the behavior of released parasites, we determined the maximum egress distance that parasites moved away from the original vacuole or cyst following egress. Tachyzoites migrated much further than bradyzoites after induced egress (**Figure 1C**). Bradyzoites also moved more slowly than tachyzoites (**Figure 1D**), as shown by quantification of their trajectories from time lapse video microscopy images. Taken together, these findings indicate that egress by bradyzoites in response to Ca^2+^ ionophore or zaprinast is incomplete and restricted.

**Figure 1.**
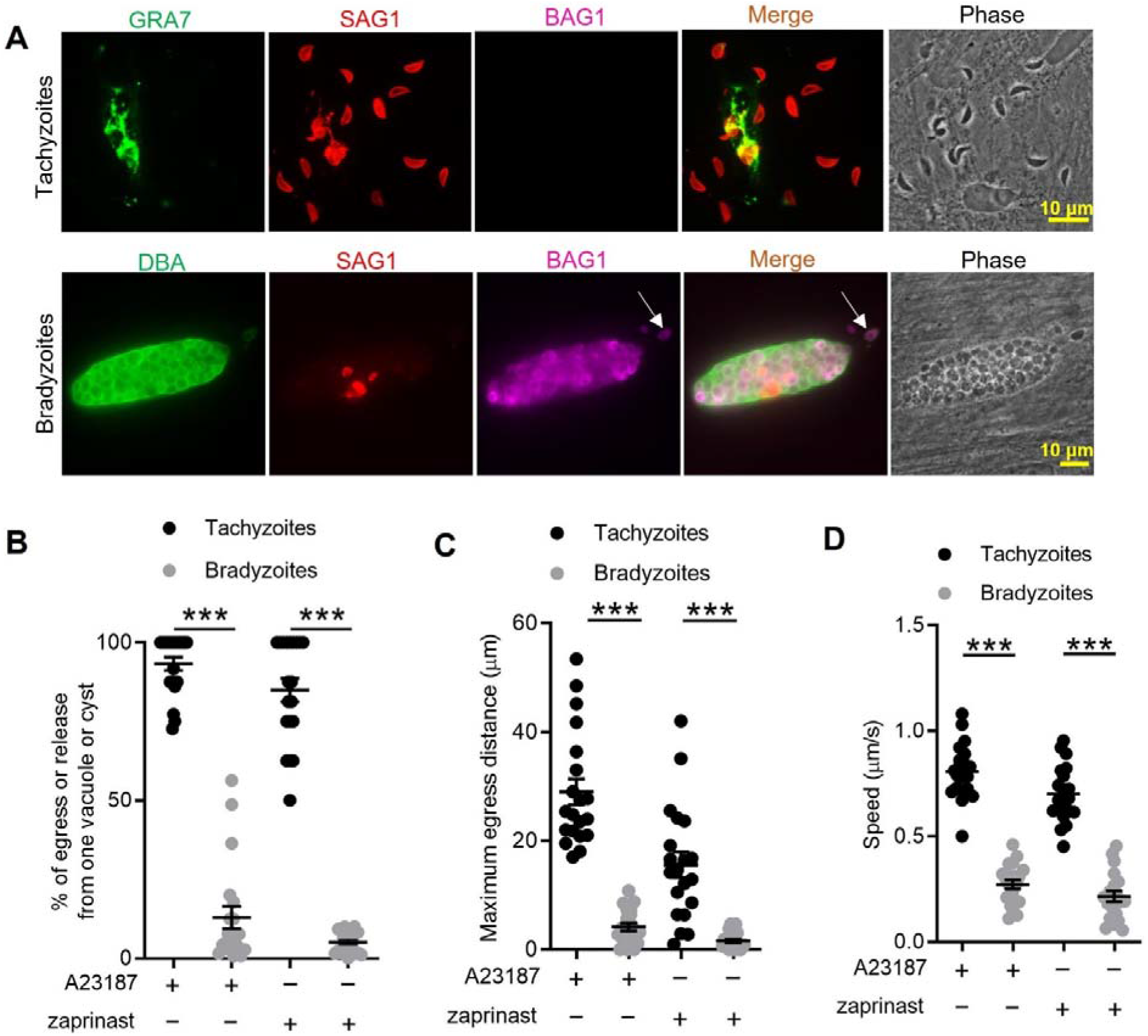
In vitro induced bradyzoites show limited egress in response to Ca^2+^ agonists. (A) Egress of tachyzoites and bradyzoites in response to A23187 (2 μM) for 15 min. Anti-GRA7, anti-SAG1, and anti-BAG1 antibodies followed by secondary antibodies to Alexa conjugated fluorochromes were used to detect the parasitophorous vacuole (PV) membrane, tachyzoites, and bradyzoites, respectively. DBA (*Dolichos biflorus* agglutinin) conjugated to FITC was used to stain the cyst wall. Arrow indicates released bradyzoites. Scale bar = 10 μm. (B) Quantitative analysis of egress in response to A23187 (2 μM) or zaprinast (500 μM) in extracellular buffer (EC) with Ca^2+^ for 15 min. Each data point represents the % of egressed or released parasites from one parasitophorous vacuole (PV) or cyst (n=20). Means ± SD of two independent experiments with 20 replicates. Two-tailed Mann-Whitney test, \*\*\**P* < 0.001. (C) Quantitative analysis of maximum distance egressed or released parasites moved away from the vacuole/cyst in response to A23187 (2 μM) or zaprinast (500 μM) in EC buffer with Ca^2+^ for 15 min. Each data point represents distance travelled of one egressed tachyzoite or released bradyzoite from the original PV or cyst (n=20). Means ± SD of two independent experiments with 20 replicates. Two-tailed Mann-Whitney test, \*\*\**P* < 0.001. (D) Quantitative analysis of speed (μm/s) of egressed or released parasites in response to A23187 (2 μM) or zaprinast (500 μM) in EC buffer with calcium for 15 min by time-lapse microscopy. Mean speed was determined by time lapse recording during the first 1 min after egress or release. Each data point represents migration speed of a single egressed tachyzoites or released bradyzoites from original PV or cyst (n=20). Means ± SD of two independent experiments with 20 replicates. Two-tailed unpaired Student’s t test, \*\*\**P* < 0.001.

### Calcium-mediated microneme secretion is dampened by bradyzoite development

Egress by parasites requires Ca^2+^-stimulated microneme secretion. To examine the reason for inefficient egress by bradyzoites, we monitored microneme secretion by quantitative secretion analysis of MIC2 fused with *Gaussia* Luciferase (Gluc). The *MIC2-Gluc* reporter was randomly integrated into the genome of the BAG1-mCherry strain (**Figure 2A**). IFA revealed that MIC2-Gluc was expressed and localized to micronemes in tachyzoites and bradyzoites induced for 7 days at pH 8.2 in vitro, as confirmed by expression of BAG1-mCherry (**Figure 2B**). BAG1-mCherry MIC2-GLuc strain tachyzoites, and bradyzoites liberated from cysts produced by cultivation for 7 days at pH 8.2 in vitro, were sorted by FACS (**Figure 2C**). FACS sorted tachyzoites and bradyzoites were treated with zaprinast or ionomycin, a Ca^2+^ ionophore that induces release of Ca^2+^ from the ER [50]. Bradyzoites secreted much less MIC2-Gluc protein compared to tachyzoites in response to Ca^2+^ agonists, zaprinast and ionomycin as shown by *Gaussia* luciferase assays performed on ESA fractions collected following stimulation (**Figure 2D**). To further investigate the process of microneme secretion by bradyzoites, we randomly integrated a mCherry secretion reporter, based on the signal peptide sequence of ferredoxin-NADP(+)-reductase (FNR-mCherry), into the genome of BAG1-EGFP parasites (**Figure 2E**). The FNR-mCherry reporter is an improved version of DsRed reporter that is secreted into the matrix of PV, and released following the discharge of PLP1 in response to Ca^2+^ agonists [15]. Then we monitored the permeabilization of PV membrane or cyst wall after stimulation with A23187 based on the diffusion of FNR-mCherry using time-lapse fluorescence video microscopy. Consistent with previous reports [51], we observed that A23187 stimulated fast leakage of FNR-mCherry from the PV surrounding tachyzoites (**Figure 2F**, top panel and **Figure 2-video 1**). However, FNR-mCherry was not released from the cyst after A23187 stimulation (**Figure 2F,**middle panel and **Figure 2-video 3**). As a control to confirm that the FNR-mCherry was indeed secreted into the lumen of the cyst matrix, we treated cysts with trypsin to release bradyzoites. Once the cyst wall was digested, the FNR-mCherry dissipated rapidly, confirming that it was present in the matrix of the cyst (**Figure 2F**, bottom panel and **Figure 2-video 2**). These data were also confirmed by plotting FNR-mCherry fluorescence intensity changes vs. time for tachyzoites vs. intact or trypsin treated cysts (**Figure 2G**). These findings demonstrate dampened microneme secretion by bradyzoites, which may explain their incomplete egress.

**Figure 2.**
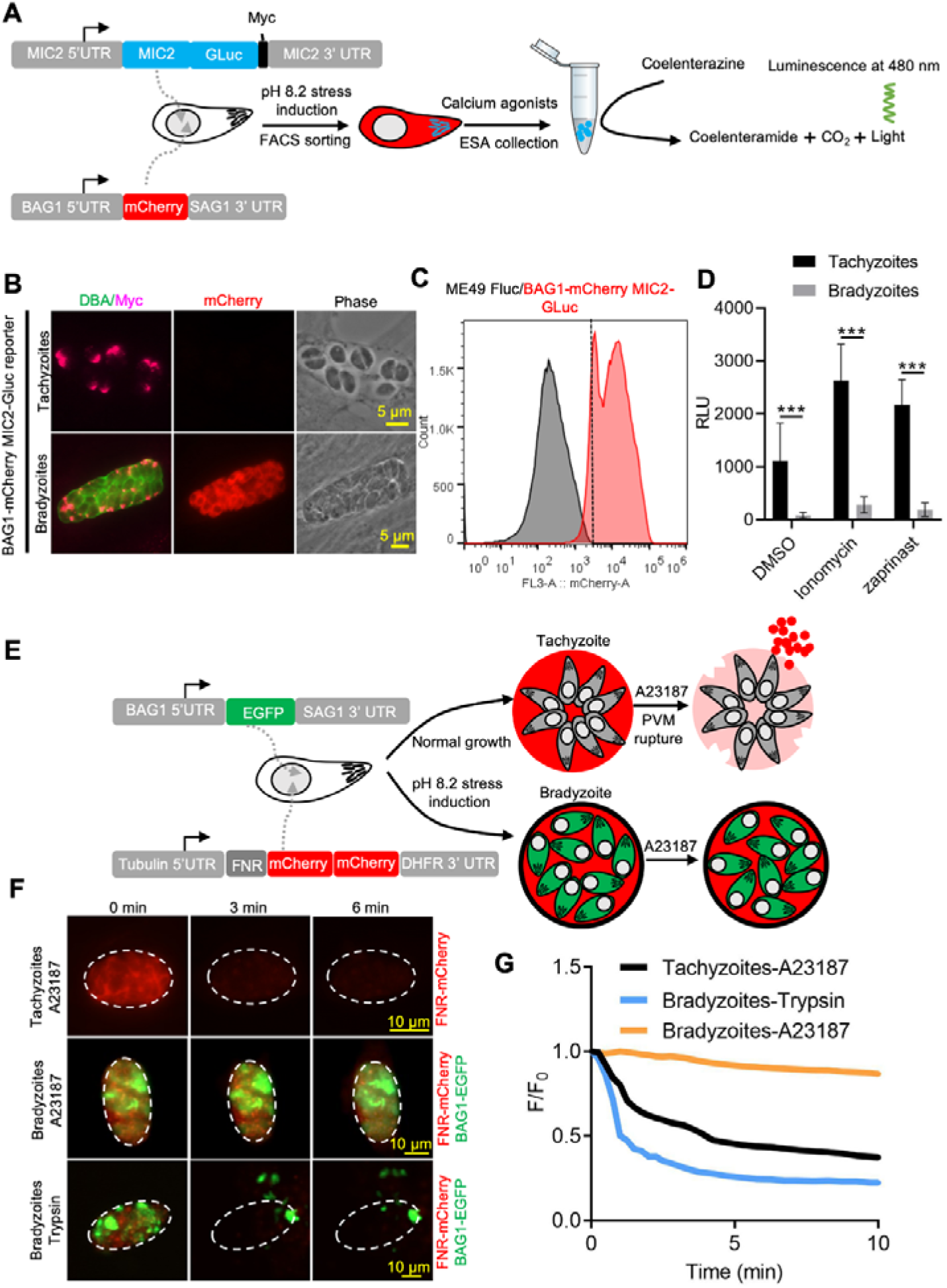
Ca^2+^ dependent microneme secretion is significantly dampened in bradyzoites. (A) Schematic of bradyzoites MIC2 secretion assay using ME49 BAG1-mCherry MIC2-GLuc bradyzoites, differentiated in vitro by cultivation at pH 8.2 for 7 days, based on fluorescence-activated cell sorting (FACS). (B) IFA analysis showing localization of MIC2-Gluc in bradyzoites induced for 7 days at pH 8.2. MIC2-Gluc was stained with anti-Myc antibody, bradyzoites were detected with anti-mCherry, followed by secondary antibodies conjugated with Alexa Fluor dyes, and the cyst wall was stained with DBA-FITC. Bar = 5 μm. (C) Bradyzoites expressing BAG1-mCherry were induced for 7 days at pH 8.2, mechanically liberated from cysts by 0.25 mg/ml trypsin for 5 min in intracellular buffer (IC buffer) and collected by FACS after gating with parental ME49 Δ*hxgprt::Fluc* parasites. (D) ME49 BAG1-mCherry MIC2-Gluc tachyzoites or bradyzoites sorted by FACS and resuspended in EC buffer with calcium were stimulated by 0.1% DMSO, ionomycin (1 μM) or zaprinast (500 μM) for 10 min at 37 °C. Release of MIC2-GLuc in ESA was determined using a *Gaussia* luciferase assay. Means ± SEM of three independent experiments each with 3 replicates. Multiple Student’s t tests, \*\*\**P* < 0.001. (E) Schematic illustration of the FNR-mCherry BAG1-EGFP dual fluorescence reporter and leakage of FNR-mCherry from the PV (top) or cyst matrix (bottom) following A23187-induced membrane permeabilization. (F) FNR-mCherry leakage was monitored by time-lapse imaging of FNR-mCherry after A23187 (2 μM) treatment. FNR-mCherry BAG1-EGFP tachyzoites cultured under normal condition for 24 hr or bradyzoites induced for 7 days at pH 8.2 were treated with A23187 (2 μM) or 0.25 mg/ml trypsin in EC buffer with calcium for 10 min at 37◻. Dash circle indicates the region of interest (ROI) for measurement of fluorescence intensity. Bar= 10 μm. (G) FNR-mCherry fluorescence (F) over the initial signal (F_0_) vs. time from cells treated as in F. Curves are the mean data of 3 independent vacuoles or cysts. Bradyzoites treated with DMSO group was used to assess photobleaching of mCherry (grey line).

### Genetically encoded calcium reporter reveals dampened Ca^2+^ responses in bradyzoites

To investigate Ca^2+^ signaling in bradyzoites, we established a dual fluorescent reporter system containing constitutively expressed GCaMP6f and mCherry under the control of bradyzoite stage-specific promoter BAG1 (**Figure 3A**). Using this system, both tachyzoites and bradyzoites express the same levels of GCaMP6f, while only bradyzoites express mCherry, allowing specific monitoring of Ca^2+^ signals in both stages. We compared the response of BAG1-mCherry GCaMP6f reporter parasites that were grown as tachyzoites, to those induced to form bradyzoites by cultivation in HFF cells for 7 days at pH 8.2 in vitro, after treatment with Ca^2+^ ionophore A23187. A23187 induced rapid and high-level increases in GCaMP6f fluorescence in tachyzoites but delayed and much weaker responses in bradyzoites as monitored by time-lapse video microscopy (**Figure 3B, Figure 3-video 1** and **Figure 3-video 2**). To determine the effect of bradyzoite development on Ca^2+^ signaling, we treated intracellular tachyzoites, vs. bradyzoites induced by cultivation in HFF cells at pH 8.2 in vitro for 4 to 7 days, and quantified time of each tachyzoite vacuole or bradyzoite cyst to reach Ca^2+^ peak level after addition of A23187 ionophore by video microscopy. Increasing time of bradyzoites development was associated with progressively longer times to reach peak fluorescence of GCaMP6f (**Figure 3C**). Time lapse recording of GCaMP6f fluorescence intensity ratio changes (F/F_0_) showed delayed Ca^2+^ increase and lower fold changes in bradyzoites compared with tachyzoites in response to A23187 stimulation (**Figure 3D**). Zaprinast also elicited slower Ca^2+^ increases and lower fold changes in bradyzoites compared with tachyzoites even in the presence of extracellular Ca^2+^ (**Figure 3E**). To better characterize Ca^2+^ responses of bradyzoites, we performed live video imaging using spinning disc confocal microscopy to distinguish individual bradyzoites within in vitro differentiated cysts and identify motile bradyzoites within cysts by comparing consecutive images (**Figure 3F**). Motile bradyzoites were also observed to have higher GCaMP6f signals and these typically oscillated over time. In response to Ca^2+^ agonists, intracellular bradyzoites showed reduced percentages of motility compared to tachyzoites (**Figure 3G**). In summary, Ca^2+^ dynamics are delayed and reduced in bradyzoites in response to Ca^2+^ agonists.

**Figure 3.**
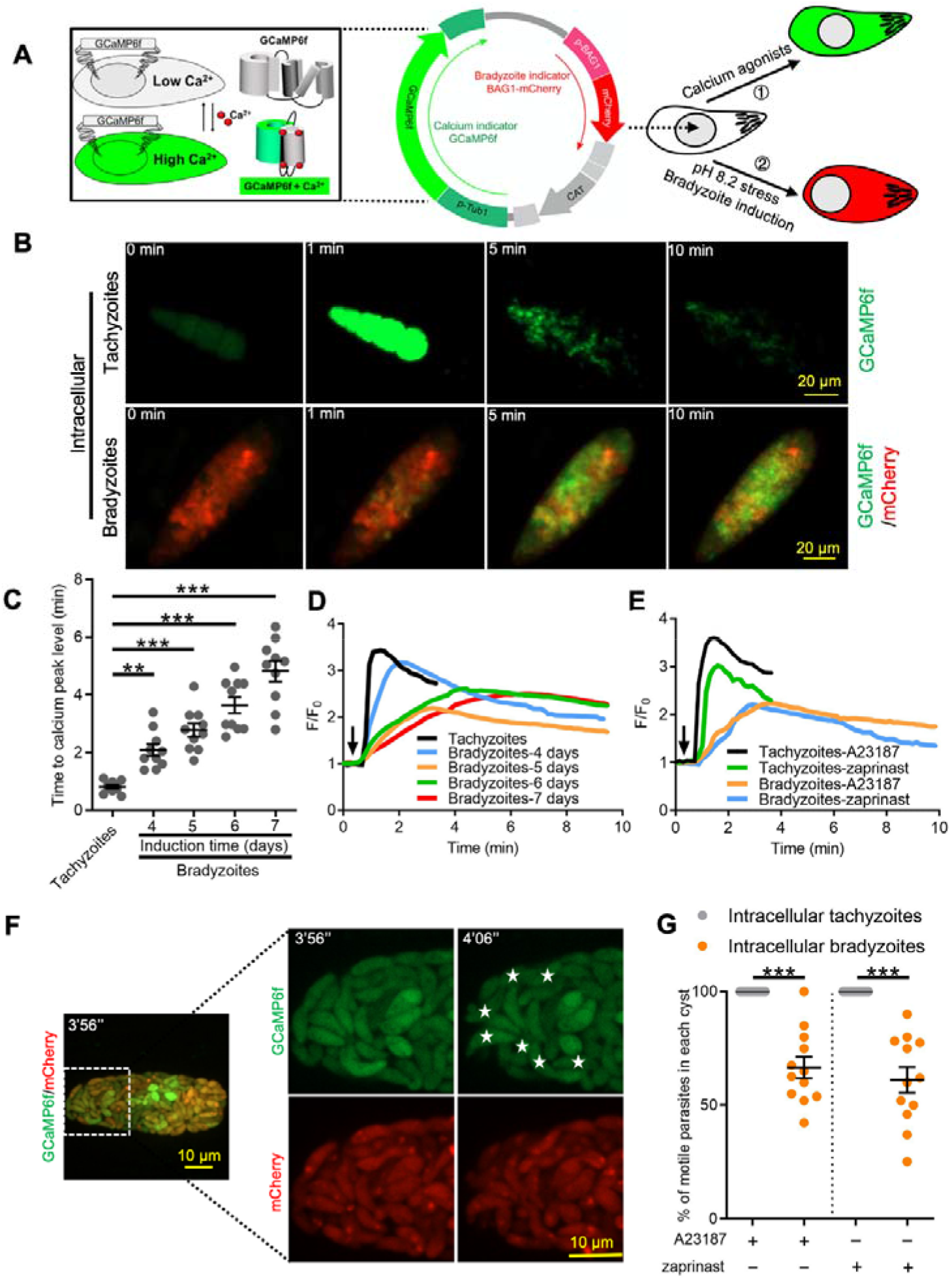
Ca^2+^ signaling is dampened during in vitro bradyzoite development induced by alkaline pH. (A) Schematic of generation of BAG1-mCherry and GCaMP6f dual fluorescent reporter to monitor Ca^2+^ responses in bradyzoites. (B) Time-lapse images BAG1-mCherry GCaMP6f tachyzoites cultured for 24 hr vs. bradyzoites induced for 7 days at pH 8.2 in response to A23187 (2 μM) in EC buffer with Ca^2+^ for 10 min. Bar= 20 μm. (C) Time for reaching Ca^2+^ peak level in response to A23187 (2 μM) for BAG1-mCherry GCaMP6f expressing tachyzoites and bradyzoites induced at pH 8.2. Data points of each group represent 10 cysts or vacuoles. Means ± SD of two independent experiments with 10 replicates each. One way ANOVA with Dunn’s multiple comparison correction test **, *P* < 0.01, ***, *P* < 0.001. (D) Monitoring the relative intensity of GCaMP fluorescence fold change (F/F_0_) vs. time for intracellular tachyzoites and in vitro induced bradyzoites induced at pH 8.2. Cells were treated with A23187 (2 μM) in EC buffer without Ca^2+^ for 10 min. Curves are the mean fluorescence intensity of 3 vacuoles or cysts. Arrow indicates time of addition of A23187. (E) Monitoring the relative intensity of GCaMP fluorescence vs. time for intracellular tachyzoites and in vitro induced bradyzoites (5 days at pH 8.2). Cells were treated with A23187 (2 μM) or zaprinast (500 μM) in EC buffer with Ca^2+^. Arrow indicates time of addition of agonists. Curves represent the mean data of 3 independent cysts or vacuoles. (F) Live time-lapse imaging of BAG1-mCherry GCaMP6f bradyzoites induced for 7 days at pH 8.2 in response to A23187 (2 μM) in EC buffer with calcium. Cells were imaged by spinning disc confocal microscopy after reaching calcium peak levels (left panel). Right panel showed its corresponding zoomed-in images. The interval between two continuous images is 10 s, white asterisks in the latter image (4’06’’) indicate motile bradyzoites by comparison with the former image (3’56’’). Bar= 10 μm. (G) Motility of parasites within PVs or cysts was analyzed by time-lapse spinning disc confocal microscopy and tracking of individual parasites for 5 min after reaching Ca^2+^ peak levels in response to A23187 (2 μM) or zaprinast (500 μM) in EC buffer with calcium. Each data point represents parasites from one vacuole or cyst (n=10). Data come from two independent experiments. Two-tailed Mann-Whitney test, \*\*\**P* < 0.001. Lines and error bars represent means ± SD of two independent experiments with 10 replicates each.

### Bradyzoites formed in skeletal muscle cell and within ex vivo cysts show diminished Ca^2+^ responses

To rule out the possibility that alkaline pH stress used for differentiation resulted in lowered Ca^2+^ signals in bradyzoites, we examined Ca^2+^ signaling in bradyzoites within cysts that formed naturally in differentiated C2C12 myocytes. Differentiated myocytes stained positively for skeletal myosin, and facilitated the development of bradyzoites, as shown using the bradyzoite stage-specific protein BAG1 (**Figure 4A**). We tested Ca^2+^ responses of bradyzoites formed in muscle cells using the dual fluorescent reporter GCaMP6f BAG1-mCherry parasites in response to A23187 or zaprinast by time-lapse video recording. Time-lapse imaging showed slow increase of GCaMP6f fluorescence in response to A23187 in tissue cysts formed in C2C12 myocytes (**Figure 4B**). Both the rate of increase and the maximum amplitude of the GCaMP6f signal was much lower in bradyzoites differentiated in myocytes compared to tachyzoites cultured in undifferentiated myoblasts (**Figure 4C**). The time to reach the peak GCaMP6f fluorescence was also delayed in bradyzoites formed in C2C12 myocytes compared with tachyzoites grown in myoblasts (**Figure 4D**). Bradyzoites cultured in C2C12 myocytes show significantly lower motility in response to A23187 and zaprinast when compared with tachyzoites (**Figure 4E**).

**Figure 4.**
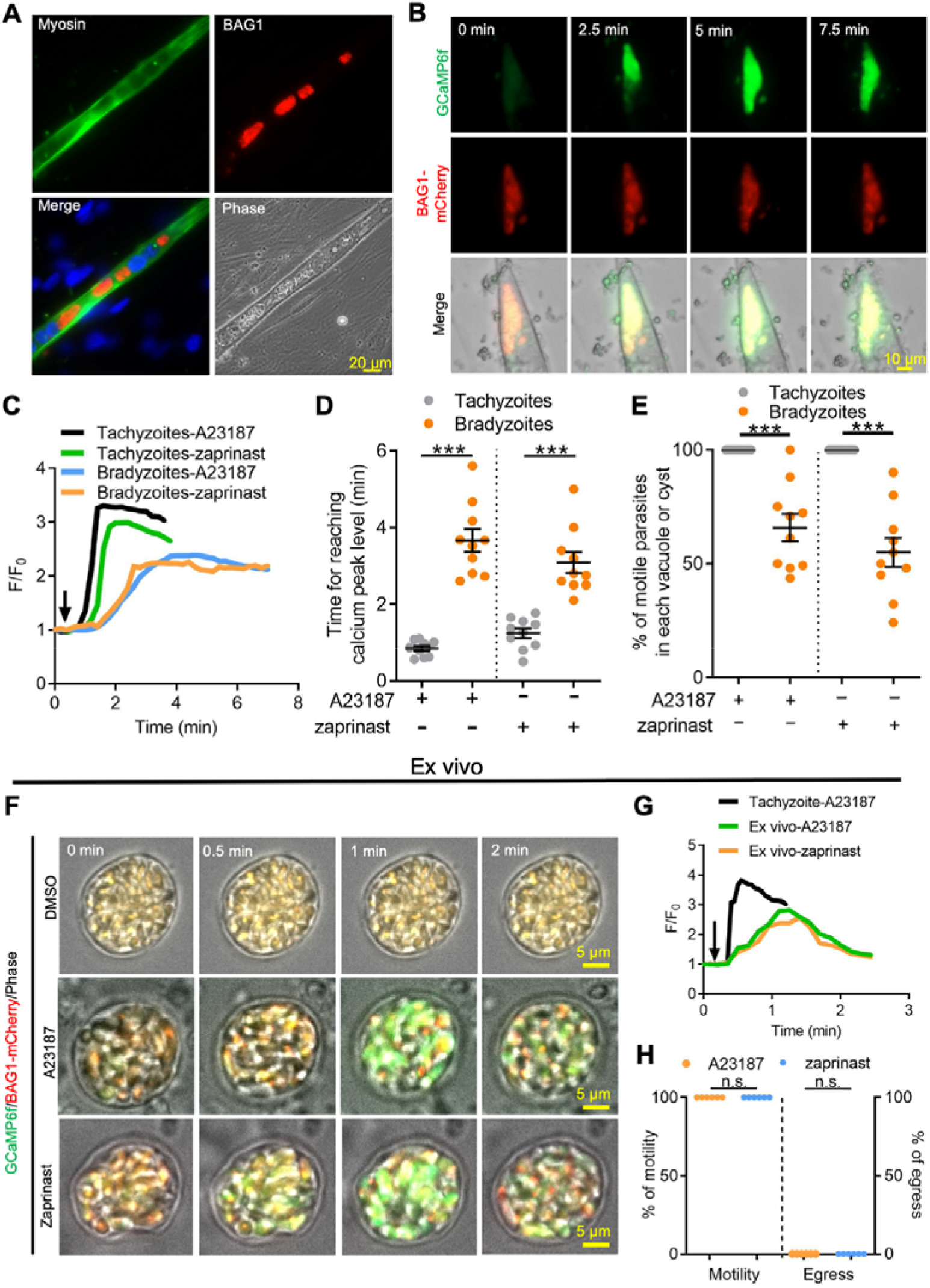
Ca^2+^ signaling is dampened in in vitro bradyzoites from spontaneously formed cysts in C2C12 muscle cells and cysts isolated from chronically infected mice. (A) Microscopy based assay for detection of bradyzoites naturally formed after 7 days culture of the BAG1-mCherry GCaMP6f expressing dual reporter strain in differentiated C2C12 muscle cells. Anti-myosin antibody was used to confirm the differentiation of C2C12 cells while BAG1 was used to detect bradyzoites followed by secondary antibodies conjugated with Alexa Fluor dyes. Bar = 20 μm. (B) Time-lapse recording of GCaMP6f fluorescence intensity from cysts of the BAG1-mCherry GCaMP6f strain naturally formed after 7 days culture in C2C12 cells. Cells were treated with A23187 (2 μM) in EC buffer with Ca^2+^. Bar = 10 μm. (C) GCaMP6f fluorescence intensity changes vs. time from tachyzoites cultured in undifferentiated myoblasts or cysts naturally formed after 10 days in differentiated C2C12 cells in response to A23187 (2 μM) or zaprinast (500 μM) in EC buffer with calcium. Curves represent mean data of 3 independent cysts or vacuoles. (D) Time for reaching Ca^2+^ peak levels in tachyzoites cultured in undifferentiated myoblasts and bradyzoites formed after 10 days culturing in C2C12 cells. Cells were treated with A23187 (2 μM) or zaprinast (500 μM) in EC buffer with calcium for 10 min. Data points of each group come from 10 cysts or vacuoles of two independent experiments. Two-tailed unpaired Student’s t test, \*\*\**P* < 0.001. Lines represent means ± SD of two independent experiments with 10 replicates each. (E) Motility of parasites analyzed by time-lapse spinning disc confocal microscopy and tracking of individual parasites for 5 min after reaching calcium peak levels in response to A23187 (2 μM) or zaprinast (500 μM) in EC buffer with calcium. Lines represent means ± SD of two independent experiments with 10 replicates each. Two-tailed Mann-Whitney t test, \*\*\**P* < 0.001. (F) Monitoring of GCaMP fluorescence in response to 0.1% DMSO, A23187 (2 μM) or zaprinast (500 μM) in EC buffer with Ca^2+^ in ex vivo cysts isolated from the brains of mice infected with BAG1-mCherry GCaMP6f reporter parasites. Cysts were harvested at 30 days post infection. Bar = 5 μm. (G) GCaMP6f fluorescence intensity changes vs. time within BAG1-mCherry GCaMP6f ex vivo cysts in response to A23187 (2 μM) or zaprinast (500 μM) in EC buffer with calcium. Curves are the mean data of 3 independent cysts. (H) Quantitative analysis of motility and egress by bradyzoites from ex vivo cysts isolated from CD-1 mice brain tissues at 30 days post-infection. Motility was analyzed by time-lapse microscopy and tracking of individual parasites using time points similar to D, E above. Each data point represents percentage of motile or egressed parasites from one cyst (n=5). Significance was determined by two-tailed Student’s t-test, n.s., not significant.

To further examine Ca^2+^ signaling in bradyzoites, we harvested tissue cysts containing BAG1-mCherry GCaMP6f bradyzoites from the brains of chronically infected CD-1 mice and investigated their responses ex vivo. Video microscopy of ex vivo tissue cysts showed slow increases in GCaMP6f fluorescence in response to A23187 or zaprinast (**Figure 4F**). The ratio of GCaMP6f fluorescence changes vs time (F/F_0_) from bradyzoites within ex vivo cysts demonstrated lower and slower changes, consistent with lower Ca^2+^ levels, compared with extracellular tachyzoites in response to Ca^2+^ agonists (**Figure 4G**). In comparing the response of extracellular, ex vivo tissue cysts (**Figure 4 F,G**) to intracellular cysts formed during infection of C2C12 myocytes (**Figure 4 B,C**), it was evident that the extracellular cysts respond somewhat faster, albeit still much slower than tachyzoites. This intermediate level of response was also seen in in vitro differentiated tissue cyst (produced by cultivation in HFF cells at pH 8.2 for 7 days) that were liberated from HFF cells and tested in vitro (**Figure 4-supplement 1**). Next, we measured the percentage of motile and egressed bradyzoites within ex vivo tissue cyst treated with A23187 and zaprinast. Strikingly, no egressed bradyzoites were observed although all the bradyzoites within ex vivo cysts became motile after stimulation (**Figure 4H**, **Figure 4-video 1**, **Figure 4-video 2**). Taken together, these findings indicate that bradyzoites formed spontaneously in muscle myocytes and within ex vivo cysts from chronically infected mice display dampened Ca^2+^ dynamics when treated with Ca^2+^ agonists.

### Bradyzoites store less Ca^2+^ in ER and acidocalcisome

The cyst wall surrounding bradyzoites may restrict access to Ca^2+^ agonists and hence dampen signals from GCaMP6f in response to Ca^2+^ agonists in the studies described above. To test this possibility, we monitored GCaMP6f fluorescence changes in extracellular bradyzoites vs. tachyzoites of the BAG1-mCherry GCaMP6f strain by live imaging. Bradyzoites were induced by cultivation in HFF cells at pH 8.2 for 7 days and liberated from cysts by trypsin treatment, followed by washing and resuspension for analysis. We also observed slower increases in GCaMP6f fluorescence intensity in bradyzoites (**Figure 5-video 2**) compared with tachyzoites (**Figure 5-video 1**) in response to A23187 (**Figure 5A**). Quantitative analysis of Ca^2+^ fluorescence changes (F/F_0_) after stimulation by A23187 and zaprinast showed slower Ca^2+^ responses in extracellular bradyzoites when compared to tachyzoites (**Figure 5B**). To confirm that extracellular bradyzoites were viable after liberation from in vitro cultured cysts by trypsin treatment, we utilized SYTOX Red, which is a DNA dye excluded by intact membranes of viable cells. In contrast to bradyzoites that were formaldehyde-fixed as a positive control, extracellular bradyzoites were not stained by SYTOX after the liberation from in vitro cysts (**Figure 5C**), indicating they were still viable after trypsin treatment.

**Figure 5.**
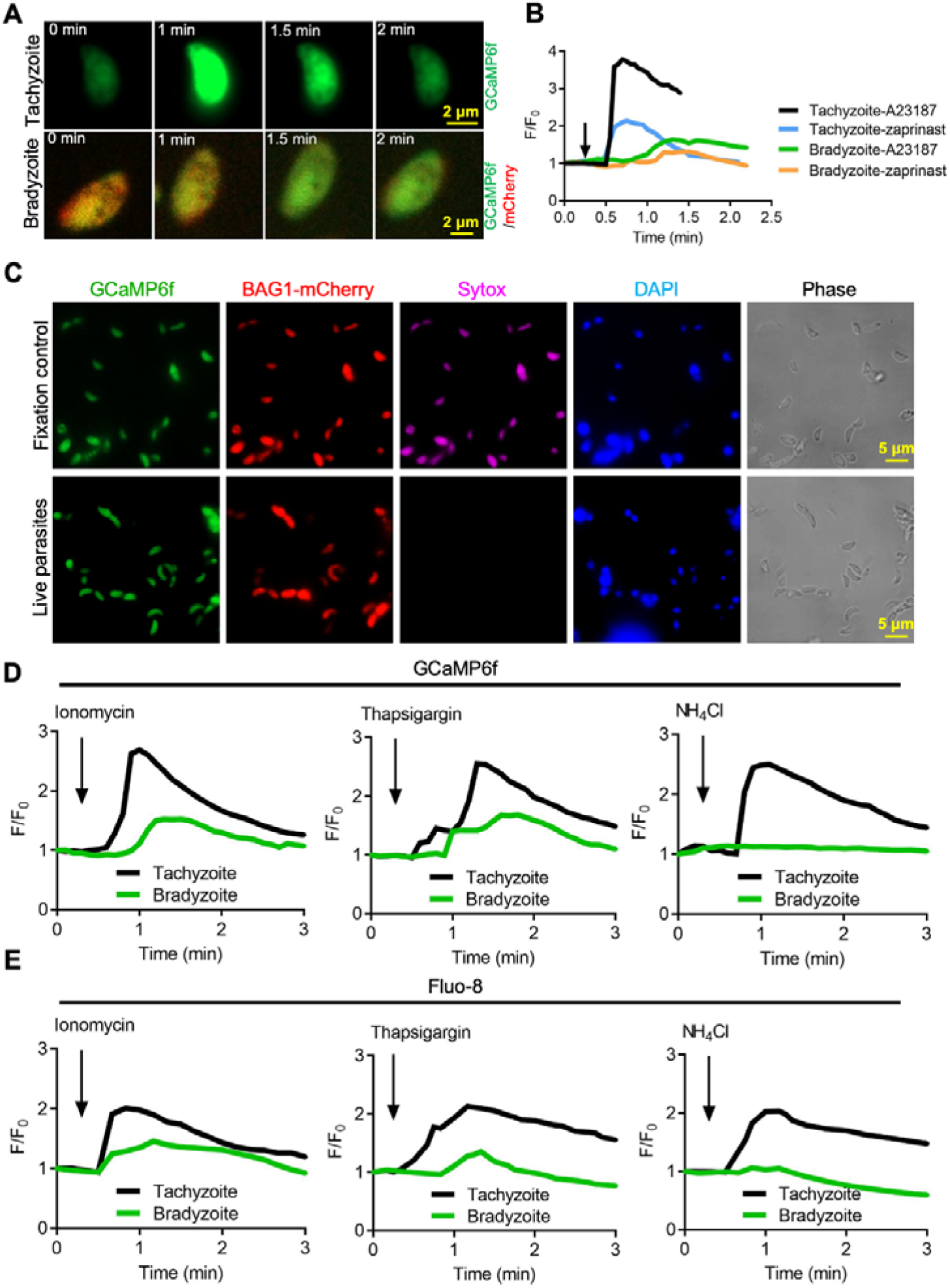
Bradyzoites have lower Ca^2+^ stores and reduced responses to agonists compared to tachyzoites. (A) Live imaging of extracellular BAG1-mCherry GCaMP6f dual fluorescent reporter tachyzoites and bradyzoites induced for 7 days at pH 8.2 in response to A23187 (2 μM) in EC buffer with Ca^2+^. Bar= 2 μm. (B) Fluorescence recording of increased GCaMP6f fluorescence with Ca^2+^ increase in response to A23187 (2 μM) or zaprinast (500 μM) in EC buffer with Ca^2+^ for extracellular tachyzoites and bradyzoites. Arrow indicates the addition of calcium agonists. Each curve is the mean of three individual parasites. (C) BAG1-mCherry GCaMP6f reporter live bradyzoites were stained by SYTOX^TM^ far red to detected dead cells and DAPI 30 min after liberation from cysts. Formaldehyde-fixed bradyzoites serve as positive control. Bar= 5 μm. (D) GCaMP6f fluorescence intensity vs. time for extracellular BAG1-mCherry GCaMP6f dual reporter parasites in response to 1 μM ionomycin, 1 μM thapsigargin, or 10 mM NH_4_Cl in EC buffer without Ca^2+^. Arrow indicates the addition of agonist. Each curve is the mean of three individual parasites. (E) Fluorescence intensities change fold vs. time of extracellular BAG1-mCherry expressing bradyzoites loaded with 500 nM Fluo-8 AM after addition of 1 μM ionomycin, 1 μM thapsigargin or 10 mM NH_4_Cl in EC buffer without Ca^2+^. Arrow indicates the addition of agonist. Each curve is the mean of three individual parasites.

We hypothesized that bradyzoites might have dampened GCaMP6f responses because they fail to release Ca^2+^ from intracellular stores. We tested Ca^2+^ responses of BAG1-mCherry and GCaMP6f -expressing bradyzoites and tachyzoites treated with ionomycin, which releases Ca^2+^ mainly from the ER [50], thapsigargin, which inhibits SERCA-type Ca^2+^-ATPase causing an increase of cytosolic Ca^2+^ due to uncompensated leakage from the ER [33], and NH_4_Cl, an alkalizing reagent that releases Ca^2+^ from acidic stores like acidocalcisomes [35]. Both ionomycin and thapsigargin induced delayed and lower amplitude changes in GCaMP6f fluorescence in bradyzoites vs. tachyzoites as shown by plotting fluorescence intensity fold changes (F/F_0_) vs. time (**Figure 5D**), indicative of lower ER stored Ca^2+^. In contrast, bradyzoites treated with NH_4_Cl showed no meaningful change in GCaMP6f fluorescence, suggesting they lack mobilizable acidic Ca^2+^ (**Figure 5D**). To rule out the possibility that the Ca^2+^ indicator GCaMP6f is less sensitive in bradyzoites due to some intrinsic defect, we loaded BAG1-mCherry expressing tachyzoite or bradyzoites with the Ca^2+^ sensitive vital dye Fluo-8 AM and used these cells for imaging. Fluo-8 AM labeled bradyzoites displayed dampened Ca^2+^ signaling after stimulation by ionomycin, thapsigargin or NH_4_Cl, relative to tachyzoites that responded normally (**Figure 5E**). Collectively, these findings indicate that bradyzoites are less able to mobilize Ca^2+^ from the ER and acidic stores in response to agonists.

### Ratiometric sensor reveals reduced basal levels of Ca2+ and dynamics in bradyzoites

To more precisely compare Ca^2+^ levels in tachyzoites and bradyzoites, we constructed a ratiometric fluorescence reporter by co-expression of GCaMP6f with blue fluorescent protein mTagBFP2 linked by a P2A split peptide (**Figure 6A, Figure 6 Supplement 1A, Figure 6 Supplement 1B**). Because both proteins are co-expressed from the same promoter, the mTagBFP2 serves as a control for expression level, as mTagBFP2 is non-responsive to Ca^2+^ levels [52]. Live fluorescence microscopy showed simultaneous expression of GCaMP6f and mTagBFP2 in tachyzoites, and additionally mCherry in bradyzoites (**Figure 6B**). Equal expression of GCaMP6f (His tag) and mTagBFP2, as well as separation of tachyzoites and bradyzoite populations (detected with SAG1 and BAG1 respectively) was validated by western blotting (**Figure 6C**). To compare Ca^2+^ basal levels, we quantified the fluorescence intensity ratio F_GCaMP6f_/F_mTagBFP2_ of intracellular and extracellular tachyzoites and bradyzoites in EC buffer with or without Ca^2+^. We observed significant reductions in the fluorescence intensity ratio of both intracellular and extracellular bradyzoites relative to tachyzoites (**Figure 6D**), indicative of lower resting Ca^2+^ levels in bradyzoites. We next compared Ca^2+^ dynamics of intracellular tachyzoites and bradyzoites in response to Ca^2+^ agonists ionomycin, NH_4_Cl and thapsigargin. Changes in the fluorescence of GCaMP6f were much slower and of lower amplitude in bradyzoites relative to tachyzoites (**Figure 6E**). We also observed lower resting Ca^2+^ and peak levels in extracellular bradyzoites compared to tachyzoites (**Figure 6F**), indicating lower activity or expression of cytoplasmic influx mechanisms like the PM entry or ER release channels. To understand the molecular basis for the reduced stored Ca^2+^ and responses in bradyzoites, we performed real-time PCR to compare mRNA expression levels of TgSERCA [34], which is the drug target of thapsigargin and transfers Ca^2+^ from the cytosol of parasites to ER, TgA1 [36], which plays important roles in the accumulation of Ca^2+^ in the acidocalcisome and other acidic stores, TgTRPPL-2 [53], which is a transient receptor potential (TRP) channel key for Ca^2+^ influx into the cytosol, and other calcium-related proteins, such as TgPMCA1, TgA2 and the Ca^2+^/H^+^ exchanger [54]. We observed significant reduction in the relative expression level of TgSERCA, TgA1, TgPMCA1, TgA2, Ca^2+^/H^+^ exchanger and TgTRPPL-2 in bradyzoites compared to tachyzoites (**Figure 6G**). Taken together, these findings indicate that bradyzoites have lower levels of stored Ca^2+^, which is associated with the overall downregulation of Ca^2+^-related pumps and channels.

**Figure 6.**
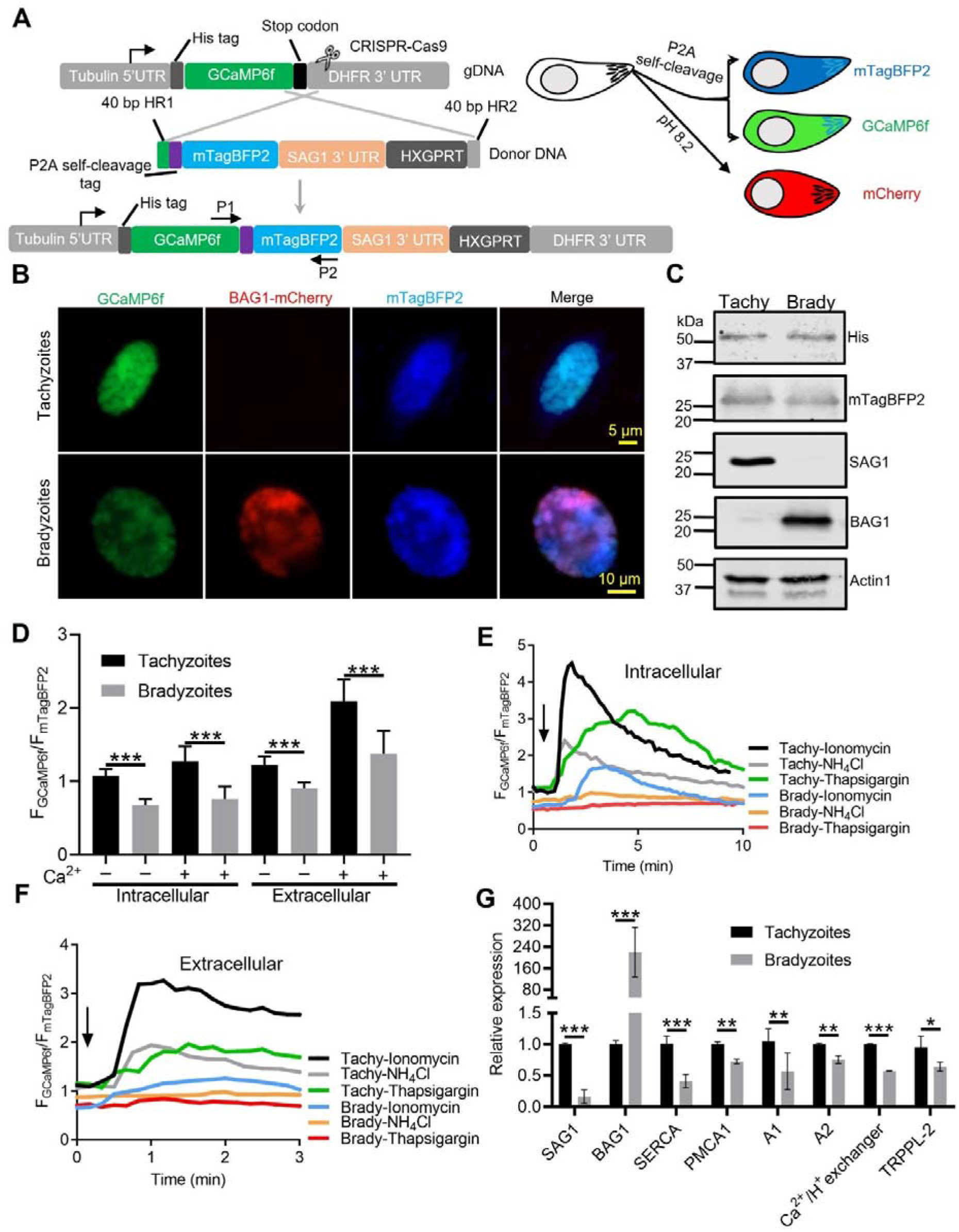
Ratiometric Ca^2+^ imaging of bradyzoites reveals lower levels of resting Ca^2+^ and reduced response to Ca^2+^ ionophores compared to tachyzoites. (A) Schematic diagram of generation of a ratiometric calcium reporter containing GCaMP6f fused with by a peptide P2A and blue fluorescence indicator mTagBFP2 in the background of BAG1-mCherry reporter strain. (B) Fluorescence microscopy imaging of intracellular ratiometric indicator expressed by tachyzoites cultured for 24 hr vs. bradyzoites induced for 7 days at pH 8.2 culture in EC buffer without Ca^2+^. Bar= 10 μm. (C) Western blots showing GCaMP6f and mTagBFP2 produced from the ratiometric reporter expressed by tachyzoites and bradyzoite. αHis and αtRFP antibodies were used to probe the expression of GCaMP6f and mTagBFP2, respectively. SAG1 and BAG1 serve as the stage-specific marker of tachyzoites and bradyzoites, respectively. Actin functions as loading control. (D) Quantification of basal calcium levels normalized by comparison of GCaMP6f to mTagBFP2 fluorescence intensity ratios of intracellular and extracellular tachyzoites or bradyzoites that were induced by culture for 7 days at pH 8.2. For extracellular parasites, tachyzoites were liberated mechanically and bradyzoites were liberated by trypsin treatment. Parasites within intact cells, or extracellular parasites were incubated in EC buffer with or without Ca^2+^ for 10 min before imaging. Data represent mean values from two independent experiments with 10 total vacuoles or cysts for each treatment. Two-tailed unpaired Student’s t test, ***, *P* < 0.001. (E) Monitoring of GCaMP6f/ mTagBFP2 fluorescence intensity ratio vs. time for intracellular tachyzoites and in vitro induced bradyzoites that were induced by culture for 7 days at pH 8.2. (F) For extracellular parasites, tachyzoites were liberated mechanically and bradyzoites were liberated by trypsin treatment. Parasites were incubated in EC buffer without Ca^2+^ for 10 min and responses were measured to ionomycin (1 μM), thapsigargin (1 μM) or 10 mM NH_4_Cl. Arrow indicates time of addition of agonists. Each kinetic curve represents mean data of 3 independent samples (individual vacuoles or cysts for intracellular and single parasites for extracellular). (G) Gene expression levels in tachyzoites and bradyzoites induced for 7 days at pH 8.2. mRNA levels were measured using RT-PCR and expressed relative to the housekeeping transcript for actin. SAG1 and BAG1 were used to monitor tachyzoites and bradyzoites, respectively. Data represent the mean ± SD of two independent assays containing triplicate samples each. Multiple Student’s t tests, **, *P* < 0.01, ***, *P* < 0.001.

### Calcium signaling plays a critical role in gliding motility of bradyzoites

To test whether dampened Ca^2+^ signaling would still be sufficient to drive gliding motility of bradyzoites, we treated BAG1-mCherry GCaMP6f expressing cysts cultured in vitro with trypsin to liberate bradyzoites (**Figure 7A**). There were no obvious changes in the Ca^2+^ levels nor motility during trypsin treatment and release (**Figure 7B** and **Figure 7-video 1**). When we monitored the motility of released bradyzoites by time-lapse video microscopy, a number of bradyzoites underwent circular gliding (**Figure 7C** and **Figure 7-video 2**) in patterns that were highly reminiscent of tachyzoite motility. Similar to previous descriptions of oscillating Ca^2+^ patterns in gliding tachyzoites [39], we observed fluctuations of GCaMP6f fluorescence intensities in single extracellular bradyzoites with gliding motility (**Figure 7D**).

**Figure 7.**
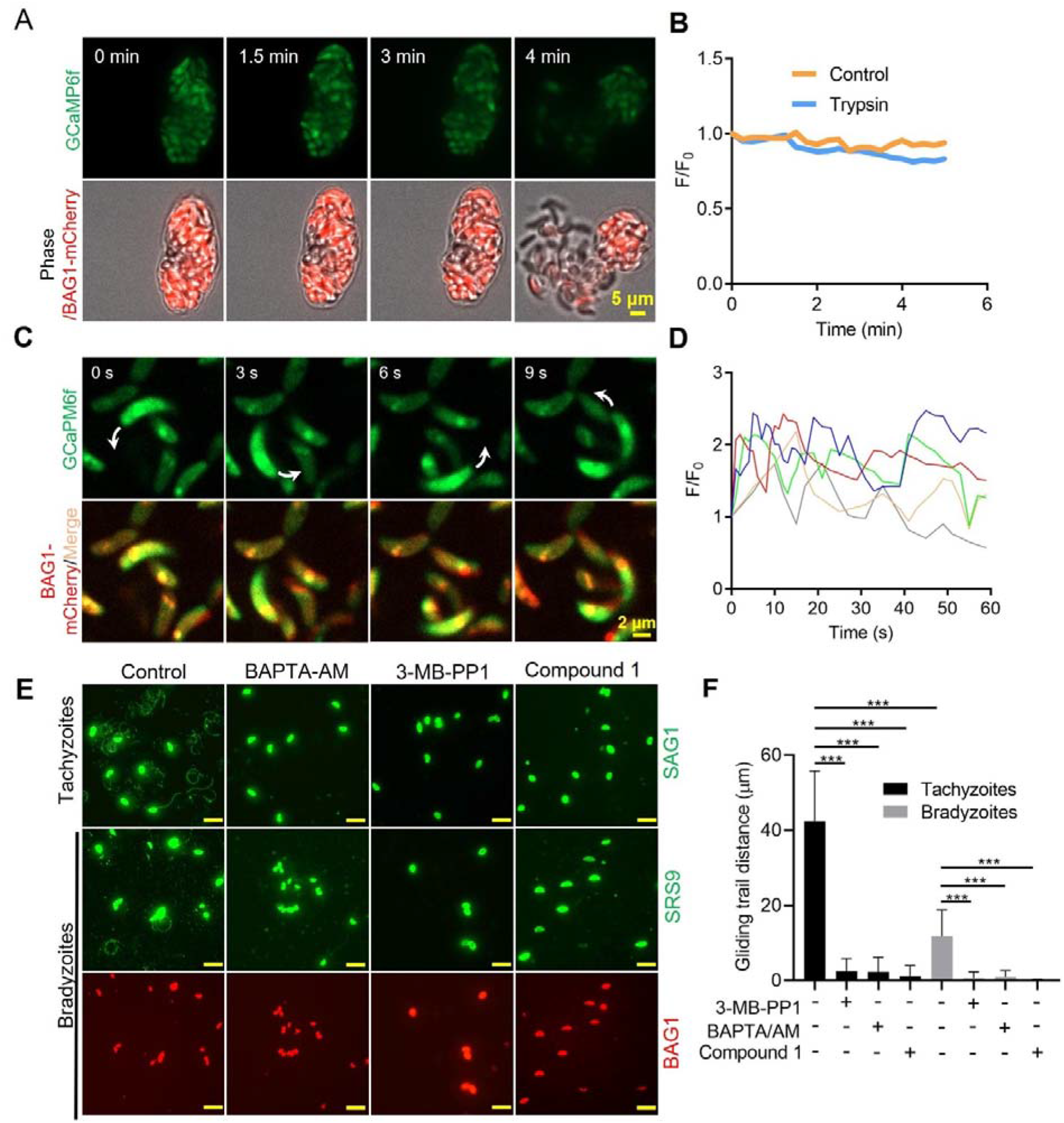
Ca^2+^ signaling governs gliding motility of bradyzoites. **(A)** Time-lapse microscopy recording GCaMP6f BAG1-mCherry bradyzoites induced for 7 days at pH 8.2. Cells were imaged during the digestion by 0.25 mg/ml Trypsin for 5 min in EC buffer with 1.8 mM Ca^2+^. Bar = 5 μm. (B) GCaMP6f fluorescence change ratio vs. time of BAG1-mCherry GCaMP6f bradyzoites induced for 7 days at pH 8.2 treated with or without trypsin. Curves represent mean data from 3 independent cysts. (C) Spinning disc confocal microscopy monitoring circular gliding motility of bradyzoites liberated by 0.25 mg/ml trypsin for 10 min from cysts induced for 7 days at pH 8.2. Arrow shows the direction of gliding motility by one bradyzoite. Bar = 5 μm. (D) Ca^2+^ kinetics of bradyzoites undergoing gliding motility after liberation from cysts induced for 7 days at pH 8.2. The graph shows fluctuated Ca^2+^ kinetics of 5 independent single bradyzoites. (E) Indirect immunofluorescence microscopy showing the trails of parasites during gliding motility. Parasites were treated with DMSO (control), 5 μM 3-MB-PP1, 25 μM BAPTA-AM and 4 μM Compound 1. Anti-SAG1 mAb DG52 and rabbit polyclonal anti-SRS9 antibodies followed by secondary antibodies conjugated to goat anti-mouse IgG Alexa 488 were used to stain the gliding trails of tachyzoites and bradyzoites, respectively. Anti-BAG1 followed by goat anti-rabbit IgG conjugated of Alexa 568 served as marker of bradyzoites. Bar=10 μm. (F) Quantification of trails from gliding motility of tachyzoites and bradyzoites treated with DMSO (control), 5 μM 3-MB-PP1, 25 μM BAPTA-AM and 4 μM compound 1. Data represented as means ± SEM ((n = 20 replicates combined from n = 3 independent experiments). Kruskal-Wallis test with Dun’s multiple comparison correction ***, *P* < 0.001.

To further characterize the role of Ca^2+^ signaling in bradyzoites motility, we treated cells with the Ca^2+^ chelator BAPTA-AM, the PKG inhibitor compound 1, and the CDPK1 inhibitor 3-MB-PP1 to block Ca^2+^ signaling in bradyzoites. All these inhibitors significantly impaired gliding motility of tachyzoites and bradyzoites (**Figure 7E** and **Figure 7F**), confirming a key role of Ca^2+^ signaling in parasite motility. Bradyzoites displayed shorter gliding distance compared with tachyzoites as determined by measurements of trail lengths detected with SAG1 (tachyzoite) or SRS9 (bradyzoites) (**Figure 7F**), In summary, despite having dampened Ca^2+^ stores and reduced responses to agonist when intracellular, extracellular bradyzoites require calcium signaling to activate gliding motility.

### Accumulation of calcium stores and ATP synergistically activates gliding motility by bradyzoites

Following reactivation of tissue cysts, we hypothesize that bradyzoites must replenish their Ca^2+^ and energy stores to meet the demands of cell to cell transmission. To test this idea, we released bradyzoites using trypsin treatment and then treated extracellular bradyzoites with EC buffer with or without Ca^2+^ (1.8 mM) and with or without glucose (5.6 mM) for different times and stimulated the calcium responses using ionomycin. Quantitative analysis of Ca^2+^ fluorescence changes (F/F_0_) showed that bradyzoites recovered substantial stored Ca^2+^ in the presence of exogenous Ca^2+^ and glucose for 1 hr compared to 10 min (**Figure 8A and 8B**). A more modest recovery was observed in the presence of Ca^2+^ but absence of glucose (**Figure 8B**). Next, we investigated the effect of exogenous Ca^2+^ and glucose on gliding motility by bradyzoites. We used time-lapse video microscopy to determine the percentage of extracellular bradyzoites undergoing twirling, circular and helical motility after incubation in EC buffer ± Ca^2+^ and glucose for 10 min vs 1 hr. Quantitative analysis showed that bradyzoites underwent all forms of gliding motility and substantially recovered gliding motility after incubation with EC buffer containing both Ca^2+^ and glucose for 1 hr, while very few bradyzoites were able to glide following incubation with exogenous Ca^2+^ or glucose alone (**Figure 8C**).

**Figure 8.**
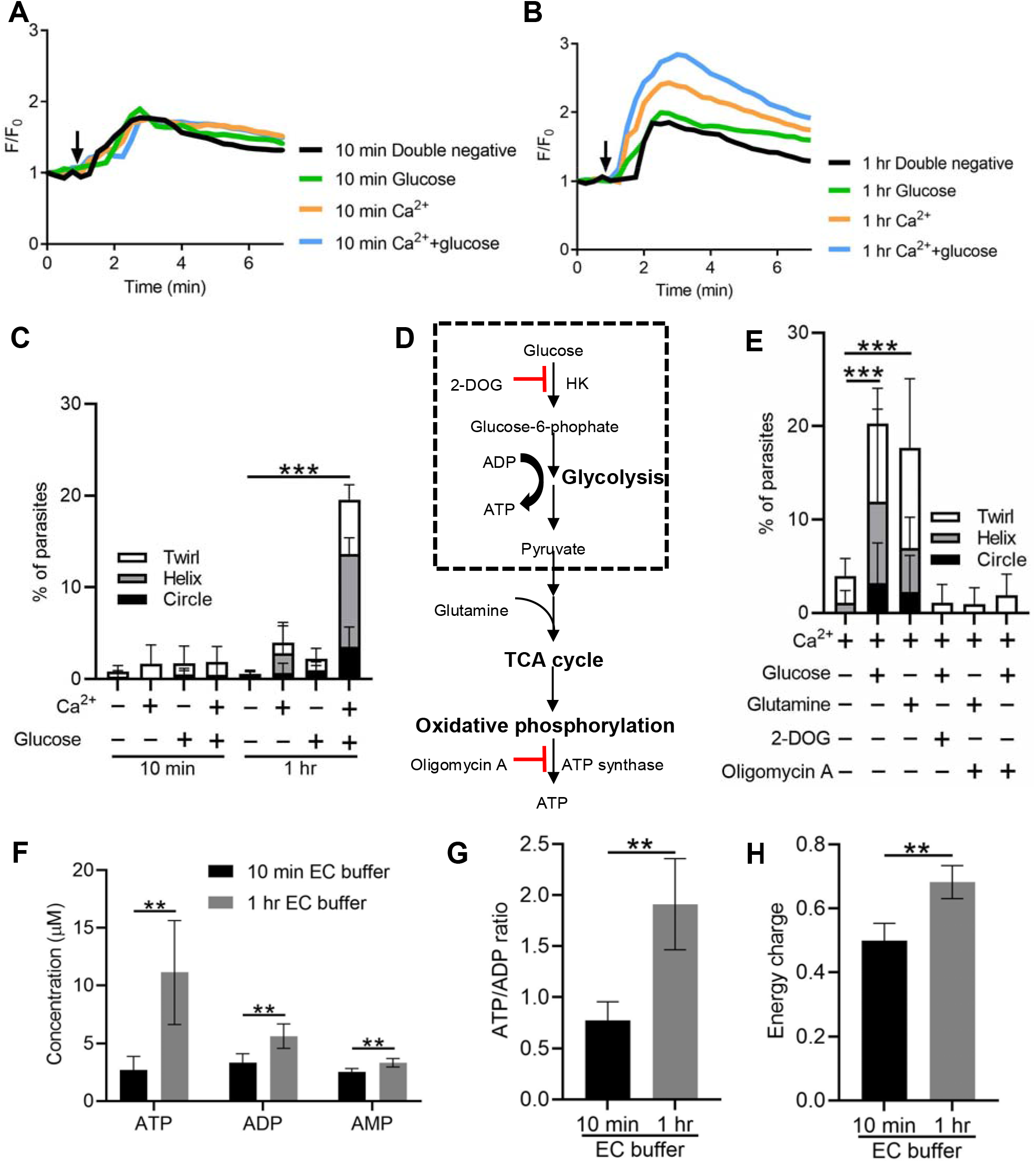
Exogenous Ca^2+^ and glucose collectively contributes to bradyzoites gliding motility via refilling calcium pools and increasing ATP production. (A-B) Monitoring the relative intensity of GCaMP6f fluorescence fold change (F/F_0_) vs. time from extracellular bradyzoites treated with 1 μM ionomycin. Bradyzoites induced for 7 days at pH 8.2 were released from in vitro cysts by 0.25 mg/ml trypsin and pre-incubated in EC buffer ± 1.8 mM Ca^2+^ and /or ± 5.6 mM glucose for 10 min (A) or 1 hr (B) before measurements. Each kinetic curve represents mean data of 3 extracellular parasites. Arrow indicates the addition of 1 μM ionomycin. Double negative refers to the absence of Ca^2+^ and glucose. (C) Percentage of extracellular parasites undergoing different forms of gliding motility as determined from time-lapse video microscopy. Bradyzoites induced for 7 days at pH 8.2 were treated in EC buffer ± 1.8 mM Ca^2+^and/or ± 5.6 mM glucose for 10 min or 1 hr before measurements. Means ± SD of two independent experiments with 6 replicates each. Kruskal-Wallis test with Dunn’s multiple comparison correction test ***, *P* < 0.001 for comparison between – calcium / - glucose and + calcium / + glucose. All other groups were not significantly different from the negative control. (D) Schematic illustration of mechanism of 2-deoxyglucose (2-DOG) and oligomycin A in inhibiting ATP production. (E) Percentage of bradyzoites with different forms of gliding motility determined by time-lapse video microscopy. Bradyzoites induced for 7 days at pH 8.2 were treated in EC buffer (1.8 mM Ca^2+^) ± 5.6 mM glucose, 5.6 mM glutamine, 50 mM 2-DOG, or 20 μM oligomycin A for 1 hr before measurements. Means ± SD of two independent experiments with 6 replicates each. Kruskal-Wallis test with Dunn’s multiple comparison correction test ***, *P* < 0.001. (F-H) High-performance liquid chromatography UV (HPLC-UV) analysis of ATP, ADP and AMP levels in extracellular bradyzoites incubated with EC buffer containing 1.8 mM Ca^2+^ and 5.6 mM glucose for 10 min or 1 hr. Bradyzoites induced for 7 days at pH 8.2 were purified by magnetic beads and released from in vitro cysts by 0.25 mg/ml trypsin. Data from two independent experiments with 6 technical replicates. (F) Concentrations of ATP, ADP, and AMP in extracellular bradyzoites represented as mean ± SD. Multiple Student’s t tests, **, *P* < 0.01. (G) ATP/ADP ratios in extracellular bradyzoites represented as mean ± SD. Two-tailed Mann-Whitney test, **, *P* < 0.01. (H) Energy charge of extracellular bradyzoites calculated as [ATP]+0.5x[ADP]/[ATP]+[ADP]+[AMP] represented as mean ± SD Two-tailed Mann-Whitney test, **. *P* ≤ 0.01.

We reasoned that exogenous glucose could be utilized by parasites to produce ATP via glycolysis or oxidative phosphorylation to maintain a variety of cellular functions. To investigate the ATP source for supporting gliding motility, we treated exogenous bradyzoites in EC buffer containing Ca^2+^ (1.8 mM) with glucose to support glycolysis vs. the glucose analogue 2-deoxy-D-glucose (2-DOG) to block glycolysis (**Figure 8D**). Alternatively, similar preparations of bradyzoites were incubated with glutamine to provide substrates for the tricarboxylic acid (TCA) cycle or the ATP synthase inhibitor oligomycin A to inhibit oxidative phosphorylation (**Figure 8D**). Quantitative analysis of percentage of gliding motility showed either glucose or glutamine significantly increased gliding motility by bradyzoites (**Figure 8E**), indicating that either carbon source can be used to produce ATP for maintaining gliding motility. Either 2-DOG or oligomycin A blocked gliding motility by bradyzoites even in the presence of exogenous glucose or glutamine (**Figure 8E**), demonstrating that both oxidative phosphorylation and glycolysis are ATP sources for driving gliding motility by bradyzoites.

To further investigate the energy status of bradyzoites, we utilized reversed-phase high-performance liquid chromatography (RP-HPLC) to measure the adenosine triphosphate (ATP), adenosine diphosphate (ADP) and adenosine monophosphate (AMP) levels in bradyzoites treated with EC buffer containing both Ca^2+^ (1.8 mM) and glucose (5.6 mM) for different time (**Figure 8-supplement 1A, 1B and 1C**). We observed that after the incubation in EC buffer for 1 hr, bradyzoites had significantly higher ATP, ADP and AMP levels (**Figure 8F**), demonstrating enhanced ATP production during incubation. The ATP/ADP ratio and energy charge have been widely used to evaluate cellular energy status, which controls the free-energy change for ATP hydrolysis for different cellular functions [55]. Bradyzoites incubated with EC buffer for 1 hr displayed significantly increased ATP/ADP ratio and energy charge (**Figure 8G and 8H**), indicating bradyzoites rapidly recover their energy status following incubation with glucose. Collectively, exogenous Ca^2+^ and glucose altogether activate bradyzoite gliding motility via restoration of ATP levels and Ca^2+^stores.

## Discussion

Calcium signaling plays important roles in the control of microneme secretion, gliding motility, and egress of apicomplexan parasites and these pathways have been extensively characterized in the tachyzoite stage of *T. gondii* [8,30], although not widely explored in other motile life cycle stages. Here we compared the responses of *T. gondii* tachyzoites and bradyzoites to Ca^2+^ ionophores and agonists that cause release of Ca^2+^ from intracellular stores and found that Ca^2+^ responses, microneme secretion, and egress by bradyzoites were all highly attenuated. Dampened Ca^2+^ responses were evident in the responses of in vitro cysts differentiated under stress conditions, naturally occurring cysts formed in muscle cells, and tissue cysts purified from brains of chronically infected mice and tested ex vivo. Reduced responses were not simply a consequence of the intracellular environment, as similar dampened Ca^2+^signals and microneme secretion were observed in single, extracellular bradyzoites. Ratiometric Ca^2+^ imaging revealed lower resting Ca^2+^ levels and reduced ER and acidic stored Ca^2+^ in bradyzoites, which is likely a reflection of down-regulation of Ca^2+^ -ATPases involved in maintaining these stores replenished. Tissue cysts are characterized by a thick wall comprised of proteins and carbohydrates which may collectively impede signals and/or restrict egress mechanically. However, when cysts were digested by trypsin to release bradyzoites, they exhibited Ca^2+^-dependent gliding motility that was enhanced by incubation in extracellular Ca^2+^ in combination with glucose, demonstrating that they express a conserved mechanism for Ca^2+^ mediated motility, albeit dampened by reduced stored Ca^2+^ and diminished energy levels. The dampened Ca^2+^ signaling responses of bradyzoites reflect adaptations that are well suited to the long-term intracellular lifestyle of these chronic stages. As well, bradyzoites retain the potential to rapidly become motile once provided with sources of energy and calcium, demonstrating remarkable physiological flexibility that favors transmission.

Egress is a crucial step in the lytic cycle of apicomplexan parasites and this response requires the sequential steps of increase in cytoplasmic Ca^2+^, secretion of micronemes, PV rupture, and activation of motility [56,57]. Our studies demonstrate that bradyzoites show minimal egress from in vitro differentiated cysts in response to agonists that normally trigger this response in tachyzoites (i.e. Ca^2+^ ionophores and zaprinast). We also demonstrate that bradyzoites are refractory to stimulation of microneme secretion using either an intracellular reporter monitoring the release of PLP1 based on the dispersion of FNR-mCherry from the cyst matrix, or a MIC2-GLuc reporter detecting secretion from extracellular bradyzoites. To explore the basis for these differences, we utilized a dual fluorescent reporter GCaMP6f BAG1-mCherry to monitor changes of cytosolic Ca^2+^ levels in bradyzoites. Calcium signaling was significantly dampened in bradyzoites as reflected in delayed Ca^2+^ spikes and lower magnitude of cytosolic Ca^2+^ increases in response to Ca^2+^ agonists. Reduced Ca^2+^ responses were also confirmed using bradyzoites naturally formed in C2C12 skeletal muscle cells and ex vivo cysts isolated from chronically infected mice, indicating that the dampened responses are not simply a consequence of alkaline pH stress during bradyzoites development in vitro. Additionally, we observed similar dampened responses from extracellular bradyzoites, indicating that decreased responses are not simply due to reduced permeability of intact cysts to agonists. To confirm these results, we also utilized Fluo-8/AM to monitor intracellular Ca^2+^ stores of bradyzoites and observed similar dampened responses. Finally, since Ca^2+^-dependent fluorescence responses by GCaMP6f or Fluo-8 are only relative and subject to differences in protein or probe levels, we developed a ratiometric calcium reporter that contains GCaMP6f fused with self-cleavage tag P2A linked mTagBFP2 under the control of the same promoter. Ratiometric measurements of the GCaMP6f signal compared to the Ca^2+^ insensitive indicator mTagBFP2, determined that bradyzoites have lower resting Ca^2+^ levels and quantitatively decreased Ca^2+^ responses relative to tachyzoites in response to Ca^2+^ agonists. Collectively, these findings conclusively show that bradyzoites have reduced Ca^2+^ responses whether developed in vitro or in vivo and using a variety of independent methods to assess both Ca^2+^ levels and physiological responses.

Based on the above findings, it seems likely that bradyzoites possess different mechanisms to control Ca^2+^ homeostasis, including differences in expression of Ca^2+^ channels and Ca^2+^ pumps relative to tachyzoites. These differences would impact Ca^2+^ storage pools, affecting cytosolic Ca^2+^ and signaling. For example, our findings indicate that bradyzoites show reduced responses to ionomycin and thapsigargin, which release Ca^2+^ from the ER, and in response to NH_4_Cl, which releases Ca^2+^ from acidocalcisomes and likely other acidic stores [35,58]. Consistent with these dampened responses, bradyzoites showed significantly reduced expression of the Ca^2+^-ATPases TgSERCA [34] and TgA1 [36], which are involved in transporting cytosolic Ca^2+^ into the ER and acidocalcisome, respectively. They also showed reduced expression of TgA2, the Ca^2+/^H^+^ exchanger and the recently described TRPPL-2 [53], which is a transient receptor potential (TRP) channel key for cytosolic Ca^2+^ influx through the plasma and ER membranes. The reduced expression of these genes is also supported by prior data on stage-specific transcriptional differences (http://Toxodb.org). Additionally, it is possible that the reduced levels of Ca^2+^ in bradyzoites reflect limitations on the availability of Ca^2+^ from the host cell, since prior studies have shown that tachyzoites acquire their intracellular Ca^2+^ from this source [37]. Further studies will be needed to decipher the contribution of these various mechanism to altered calcium homeostasis and signaling in bradyzoites.

Bradyzoites are surrounded by a cyst wall that is comprised of an outer thin compact layer and an inner sponge-like layer that faces the cyst matrix [59]. The cyst wall is enriched in dense granule proteins [60], stage-specific glycoproteins such as CST1 [61,62], and partially characterized carbohydrates [63]. This architecture may create a barrier to egress since bradyzoites were able to activate motility but not to efficiently emerge from intact cysts. We utilized trypsin to digest the cyst wall, mimicking the cyst rupture observed in chronically infected mice or following oral ingestion and exposure to pepsin [43,64]. Notably, proteolytic release did not result in immediate changes in Ca^2+^ nor motility in the parasite, suggesting that cyst wall degradation does not trigger a process akin to egress in tachyzoites. Rather, when artificially released in this manner, a subset of bradyzoites spontaneously underwent gliding motility associated with Ca^2+^ oscillations that were similar to those previously described for tachyzoites [39]. When incubated with extracellular Ca^2+^, the percentage of motile bradyzoites increased dramatically, suggesting that Ca^2+^ entry stimulates motility, similar to tachyzoites [14,41]. Unlike a previous report showing that tachyzoites contain sufficient calcium stores and energy levels to be independent of external carbon sources during the first hr after liberation [65], we observed that bradyzoites require an external source of carbon to regain Ca^2+^ stores and ATP levels. Similar to previous findings that *T. gondii* tachyzoites can support motility either from glucose through glycolysis or from glutamine that feeds into the TCA cycle [66,67], we observed that either carbon source was capable of synergizing with Ca^2+^ to restore bradyzoite motility, although inhibitor studies indicate that oxidative phosphorylation is required to restore optimal energy levels. Consistent with this prediction, we observed that bradyzoites have intrinsically low ATP/ADP ratios but that they recovered substantially when incubated extracellularly for 1 hr in Ca^2+^ and glucose. Hence, reduced expression of Ca^2+^ channels that allow influx into the cytosol and reduced expression of Ca^2+^ pumps that fill intracellular stores would result in a general reduction of stored Ca^2+^. Reduced ER Ca^2+^ could impact mitochondrial Ca^2+^, since it has been shown in mammalian cells that Ca^2+^ can be transferred directly (through membrane contact sites) from the ER to the mitochondria [68,69], which is essential for oxidative phosphorylation and ATP production. Ultimately, reduced ER Ca^2+^may be responsible for altering energy metabolism and inducing the quiescent state in *T. gondii* bradyzoites. Collectively, these findings indicate that bradyzoites are characterized by both low calcium stores and low ATP levels, but that they respond rapidly to changes in the extracellular environment to restore both energy levels and Ca^2+^ signaling systems needed for motility. Stimulation of Ca^2+^ signaling is also important in breaking dormancy [70] and pollen germination in plants [71], and initiation of the cell cycle in animal cells [72], demonstrating the important role played by Ca^2+^ signaling in reactivation.

Reduced Ca^2+^ storage, dampened Ca^2+^ signaling, and a lower energy state may reflect the long-term sessile nature of the intracellular cyst, which prolong chronic infection. The mechanisms inducing cyst wall turnover in vivo are unclear, although host cell macrophages may contribute to this process as they secrete chitinase that can lyse cysts in vitro [73]. Additionally, cyst wall turnover may be controlled by release of parasite hydrolases as suggested by the presence of GRA56, which is predicted to belong to the melibiase family of polysaccharide degrading enzymes, on the cyst wall [74]. Our in vitro studies suggest that once the cyst wall is ruptured, bradyzoites respond to higher levels of Ca^2+^ and glucose in the extracellular environment to regain motility needed for subsequent cell invasion. Emergence of bradyzoites from tissue cysts that rupture in muscle or brain, or in tissue following oral ingestion, are likely to provide an environment to recharge bradyzoites. Consistent with this idea, previous in vitro studies have shown that similar motile bradyzoites released from ruptured cysts have the ability to re-invade new host cells, establishing new cysts without an intermediate growth stage as tachyzoites [75]. Hence, the rapid metabolic recovery of otherwise quiescent bradyzoites may be important for the maintenance of chronic infection within a single host and to assure robust cellular invasion upon transmission to the next host.

## Materials and Methods

### Cell culture

*Toxoplasma gondii* tachyzoites were passaged in confluent monolayers of human foreskin fibroblasts (HFFs) obtained from the Boothroyd laboratory at Stanford University. The ME49 ∆*hxgprt::Fluc* type II strain of *T. gondii* [76] was used as a parental strain for genetic modification. Tachyzoites were cultured in Dulbecco’s modified Eagle’s medium (DMEM; Life Technologies) pH 7.4, supplemented with 10% fetal bovine serum (FBS), penicillin, and streptomycin (Life Technologies) at 37°C in 5% CO_2_. For in vitro induction of bradyzoites, parasites were cultured in alkaline medium in ambient CO_2_ as described previously [77]. In brief, infected HFF monolayers were switched to RPMI 1640 medium (MP Biomedicals) buffered to pH 8.2 with HEPES and supplemented with 5% FBS and cultured at 37°C in ambient CO_2_, during which time the alkaline medium was changed every 2 days. For spontaneous induction of bradyzoites, C2C12 muscle myoblast cells (ATCC^®^ CRL-1772™) were maintained in DMEM supplemented with 20% FBS. C2C12 myoblast differentiation and myotube formation were induced in DMEM containing 2% horse serum (Biochrom) by cultivation at 37°C in 5% CO_2_ for 5 days. Tachyzoites were inoculated into the differentiated muscle cells and cultured for another 7 days to induce bradyzoite formation, during which time the induction medium was changed every 2 days. For harvesting bradyzoites, infected monolayers were scraped into intracellular (IC) buffer (142 mM KCl, 5 mM NaCl, 1 mM MgCl_2_, 5.6 mM D-glucose, 2 mM EGTA, 25 mM HEPES, pH 7.4) and released from cells by serially passing through 18g, 20g and 25g needles, followed by centrifugation (150g, 4°C) for 10 min. The pellet containing cysts was resuspended in IC buffer. Bradyzoites were liberated from cysts by digestion with 0.25 mg/ml trypsin at room temperature for 5 min, followed by centrifugation (150g, 4°C) for 10 min. The supernatant containing liberated bradyzoites was further centrifuged (400g, 4°C) for 10 min. The pellet containing purified bradyzoites was resuspended in extracellular (EC) buffer (5 mM KCl, 142 mM NaCl, 1 mM MgCl_2_, 5.6 mM D-glucose, 25 mM HEPES, pH 7.4) with (1.8 mM Ca^2+^) or without CaCl_2_, as indicated for different assays and in the legends.

### Reagents and antibodies

A23187, zaprinast, ionomycin, thapsigargin, NH_4_Cl, Fluorescein isothiocyanate-conjugated *Dolichos biflorus* agglutinin (DBA), and BAPTA-AM were obtained from Sigma. Fluo-8 AM was obtained from Abcam. SYTOX™ Red Dead Cell Stain was obtained from Thermal Fisher. The compounds 3-MB-PP1 [51] and Compound 1 [42] were obtained as described previously. Trypsin and L-glutamine were purchased from MP Biomedicals. Adenosine 5’-triphosphate (ATP) disodium salt, adenosine 5’ -diphosphate (ADP) sodium salt, adenosine 5’ -monophosphate (AMP) disodium salt, oligomycin A and 2-deoxy-D-glucose were purchased from Sigma. Primary antibodies include mouse mAb DG52 anti-SAG1 (provided by John Boothroyd), mouse mAb 6D10 anti-MIC2 [78], rabbit anti-GRA7 [79], mouse mAb 8.25.8 anti-BAG1 (obtained from Louis Wiess), rabbit anti-BAG1 (obtained from Louis Wiess), mouse anti-c-myc (mAb 9E10, Life Technologies), mouse anti-acetylated Tubulin (mAb 6-11B-1, Sigma), rat anti-mCherry (mAb 16D7, Life Technologies), rabbit-anti SRS9 (obtained from John Boothroyd), rabbit anti-tRFP (Axxora), mouse anti-6XHis (mAbHIS.H8, Life Technologies). Secondary antibodies for immunofluorescence assays include goat anti-mouse IgG conjugated to Alexa Fluor-488, goat anti-rabbit IgG conjugated to Alexa Fluor-488, anti-mouse IgG conjugated to Alexa Fluor-568, goat anti-rat IgG conjugated to Alexa Fluor-568, goat anti-mouse IgG conjugated to Alexa Fluor-594 (Life Technologies). For Western blotting, secondary antibodies consisted of goat anti-mouse IgG, goat anti-rabbit IgG, or goat anti-rat IgG conjugated to LiCor C800 or C680 IR-dyes and detected with an Odyssey Infrared Imaging System (LI-COR Biotechnology).

### Generation of stable transgenic parasite lines

#### Dual calcium and bradyzoite reporter strain: BAG1-mCherry GCaMP6f

A dual reporter stain designed to detect bradyzoite conversion and calcium fluctuation was generated in the ME49 Δ*hxgprt::Fluc* strain [76]. We generated a plasmid named pNJ-26 that contains mCherry driven by the BAG1 promoter, the genetically encoded calcium indicator GCaMP6f under the control of Tubulin1 promoter, and selection marker cassette SAG1 promoter driving CAT. ME49 Δ*hxgprt::Fluc* tachyzoites were transfected with 20 μg pNJ-26 plasmid and selected with 20 μM chloramphenicol. Clones containing randomly integrated transgenes were confirmed by diagnostic PCR and by IFA staining. Primers are shown in Supplementary table 1.

#### Bradyzoite reporter strain: BAG1-EGFP and BAG1-mCherry

The BAG1 promoter and the mCherry open reading frame (ORF)were independently PCR amplified from pNJ-26 and the EGFP ORF was amplified from pSAG1:CAS9-U6:sgUPRT respectively. The BAG1 promoter fragment and EGFP ORF or mCherry (ORF) were cloned by NEBuilder HiFi DNA Assembly Cloning Kit (NEB, E5520S) into the vector backbone that was produced by double enzymatic digestion of pTUB1:YFP-mAID-3HA, DHFR-TS:HXGPRT using KpnI and NdeI. ME49 Δ*hxgprt::Fluc* tachyzoites were transfected with 20 μg pBAG1:EGFP, DHFFR-TS:HXGPRT or pBAG1:mCherry, DHFFR-TS:HXGPRT and selected with mycophenolic acid (MPA) (25 μg/ml) and 6-xanthine (6Xa) (50 μg/ml). Single cell clones containing randomly integrated transgenes were confirmed by diagnostic PCR and by IFA staining. Primers are shown in Supplementary table 1.

#### MIC2 secretion reporter BAG1-mCherry MIC2-GLuc

The bradyzoite reporter line BAG1-mCherry was transfected with 20 μg of the previously described pMIC2:GLuc-myc, DHFR-TS plasmid [42] and selected with 3 μM pyrimethamine (PYR). Single cell clones containing randomly integrated transgenes were confirmed by diagnostic PCR and by IFA staining.

#### FNR-mCherry leakage reporter BAG1-EGFP FNR-mCherry

The bradyzoite reporter line BAG1-EGFP was transfected with 20 μg pTUB1:FNR-mCherry, CAT (provided by the Carruthers lab) and selected with 20 μM chloramphenicol. Single cell clones containing randomly integrated transgenes were confirmed by diagnostic PCR and by IFA staining.

#### Ratiometric reporter BAG1-mCherry GCaMP6f-P2A-mTagBFP2

The ratiometric reporter strain was generated using targeted insertion with CRISPR/Cas9 using previously described methods [80] to add the blue fluorescent protein (BFP) downstream of the GCaMP6f protein in the strain BAG1-mCherry GCaMP6f. In brief, a single guide RNA (sgRNA) targeting the DHFR 3’UTR following the GCaMP6f coding sequence was generated in the plasmid pSAG1:CAS9-U6:sgUPRT [81]. The P2A-mTagBFP2 tagging plasmid was constructed by cloning a synthetic sequence containing a slit peptide (P2A) together with the blue fluorescent reporter mTagBFP2 (P2A-mTagBFP2) into the pTUB1:YFP-mAID-3HA, DHFR-TS:HXGPRT backbone by NEBuilder HIFi DNA Assembly Cloning Kit (NEB, E5520S) after double enzymatic digestion of KpnI and NdeI. Following this step, the SAG1 3’UTR was amplified from pNJ-26 and cloned into the tagging plasmid to replace DHFR 3’UTR by Gibson assembly (NEB, E5520S). BAG1-mCherry GCaMP6f reporter tachyzoites were co-transfected with 10 μg of pSAG1::CAS9-U6::sgDHFR 3’UTR and 2 μg of PCR amplified P2A-mTagBFP2-HXGPRT flanked with 40 bp homology regions, as described previously [26]. Stable transfectants were selected with 25 μg/ml MPA and 50 μg/ml 6Xa. Single cell clones containing targeted integrated transgenes were confirmed by diagnostic PCR and by IFA staining. Primers are shown in Supplementary Table S1.

### Time-lapse imaging of fluorescent reporter strains

For time-lapse microscopy, extracellular parasites were added to glass-bottom culture dishes (MatTek), or intracellular parasites were grown in host cells attached glass-bottom culture dishes. Alternating phase and fluorescent images (at different intervals specified in the legends) were collected on a Zeiss AxioObserver Z1 (Carl Zeiss, Inc.) equipped with an ORCA-ER digital camera (Hamamatsu Photonics) and a 20x EC Plan-Neofluar objective (N.A. 0.50), 37°C heating unit, and LED illumination for blue, green, red and far-red wavelengths. Spinning disk images were acquired with a 100x oil Plan-Apochromat (N.A. 1.46) objective using illumination from 488 nm and 561 nm solid state lasers (Zeiss) and Evolve 512 Delta EMCCD cameras (Photometrics) attached to the same Zeiss AxioObserver Z1 microscope. Images were acquired and analyzed using Zen software 2.6 blue edition (Zeiss). Fluorescent intensity changes (F/F_0_) vs. time were plotted with GraphPad Prism version 6 (GraphPad Software, Inc.).

### Indirect immunofluorescence assay (IFA)

Parasites grown in HFF monolayers on glass coverslips were fixed in 4% (v/v) formaldehyde in PBS for 10 min, and permeabilized by 0.25% (v/v) Triton X-100 in PBS for 20 min, and blocked in 3% bovine serum albumin (BSA) in PBS. Monolayers were incubated with different primary antibodies and visualized with secondary antibodies conjugated to Alexa Fluors. Coverslips were sealed onto slides using ProLong™ Gold Antifade containing DAPI (Thermo Fisher Scientific). Images were captured using a 63x oil Plan-Apochromat lens (N.A. 1.4) on an Axioskop2 MOT Plus Wide Field Fluorescence Microscope (Carl Zeiss, Inc). Scale bars and linear adjustments were made to images using Axiovision LE64 software (Carl Zeiss, Inc.).

### Western Blotting

Samples were prepared in 5X Laemmli buffer containing 100 mM dithiothreitol, boiled for 5 min, separated on polyacrylamide gels by SDS-PAGE, and transferred to nitrocellulose membrane. Membranes were blocked with 5% nonfat milk, probed with primary antibodies diluted in blocking buffer. Membranes were washed with PBS + 0.1% Tween 20, then incubated with goat IR dye-conjugated secondary antibodies (LI-COR Biosciences) in blocking buffer. Membranes were washed several times before scanning on a LiCor Odyssey imaging system (LI-COR Biosciences).

### Fluo-8 AM calcium monitoring

Freshly harvested parasites were loaded with 500 nM Fluo-8 AM for 10 min at room temperature, followed by centrifugation at 400 g for 5 min and washing in EC buffer without Ca^2+^. Parasites were resuspended in EC buffer without Ca^2+^ and added directly to glass-bottom culture dishes. After addition of agonists, time-lapse images were recorded and analyzed as described above.

### Egress assay

Infected cells were treated with 2 μM A23187 or 500 μM zaprinast for 15 min at 37°C. Following incubation, samples were stained by IFA using antibodies against SAG1 (mouse), GRA7 (rabbit), FITC-conjugated DBA or BAG1(rabbit) and followed by secondary antibodies conjugated to Alexa Fluors. Samples were examined by fluorescence microscopy and the percentages of egressed or released parasites per vacuole or cyst were determined at least for 20 vacuoles or cysts per experiment. The maximum egress distance of parasites from vacuole or cysts were measured from scanned tiff images in imageJ.

### Flow cytometry

ME49 BAG1-mCherry MIC2-GLuc reporter bradyzoites were induced for 7 days at pH 8.2, harvested in IC buffer as described above, and passed through 5 μm polycarbonate membrane filter. ME49 Δ*hxgprt::Fluc* tachyzoites, cultured and harvested as indicated above, were used for gating. Approximately 1 × 10^6^ parasites from each sample (ME49 BAG1-mCherry MIC2-GLuc reporter tachyzoites and ME49 BAG1-mCherry MIC2-GLuc reporter bradyzoites) were sorted on Sony SH800S Cell Sorter directly into 500 µl IC buffer followed by centrifugation. Flow cytometry data were processed using FlowJo version 10 (FLOWJO, LLC).

### Collection of excretory-secretory antigens (ESA) and Gaussia Luciferase Assay

FACS sorted MIC2-GLuc reporter tachyzoites and bradyzoites were suspended with EC buffer and incubated with different agonists at 37°C for 10 min. ESA was collected by centrifugation and mixed with Pierce™ *Gaussia* Luciferase Glow Assay Kit reagent (Thermo Scientific™) and luminescence was detected using a Cytation 3 Cell Imaging Multimode Imager (BioTek Instruments, Inc.). Buffer control values were subtracted from their corresponding sample values to correct for background.

### Real-time PCR

RNA was extracted from ME49 Δ*hxgprt::Fluc* tachyzoites and bradyzoites induced for 7 days at pH 8.2 using RNeasy Mini Kit (Qiagen) combined with QIAshredder (Qiagen) followed by DNA Removal using DNA-free™ DNA Removal Kit (Thermo Fisher) and subsequent reverse transcription using High-Capacity cDNA Reverse Transcription Kit (Thermo Fisher). Quantitative real-time PCR was performed on Applied Biosystems QuantStudio 3 Real-Time PCR System (Thermo Fisher) using SYBR^®^ Green JumpStart™ Taq ReadyMix™ (Sigma) with primers shown in Supplementary table 1. Mean fold changes from two independent experiments were calculated from ΔΔ Ct values using actin1 transcript as housekeeping gene, as described previously [82].

### Gliding trail assay

Coverslips were precoated by incubation in 50% fetal bovine serum diluted in PBS for 1 h at 37°C followed by rinsing in PBS. Freshly harvested tachyzoites or bradyzoites were resuspended in EC buffer, treated with DMSO (0.1%, v/v), or inhibitors (in 0.1% DMSO, v/v) and then added to pre-coated glass coverslips and incubated at 37°C for 15 min. Coverslips were fixed in 2.5% formalin in PBS for 10 min and the surface proteins were detected by IFA as above described using anti-SAG1 and anti-SRS9 antibodies as stage-specific markers for tachyzoites and bradyzoites, respectively. Gliding trails were captured by IFA microscopy as described above and the frequency of trails measured from tiff images using ImageJ.

### Gliding motility assay based on time-lapse video microscopy

BAG1-mCherry parasites were induced to form bradyzoites by culture at pH 8.2 in RPMI 1640 medium under ambient air (low CO_2_) for 7 days followed by scraping into IC buffer (without glucose) and repeated passage through a 23g needle. Intact, but extracellular cysts, were pellet by centrifugation at 150 *g* for 10 min and resuspended in IC buffer without glucose. During purification, all procedures were performed at 16°C. MatTek 25 mm dishes glass bottom dishes (coverslip dishes) were pre-coated with 2 ml 50% FBS at 4°C overnight and rinsed twice using PBS prior to use. Purified cysts were added to the precoated coverslip dishes in IC buffer containing 0.25mg/ml trypsin and incubated for10 min at 16°C. The medium was removed and 2 ml EC buffer ± 1.8 mM Ca^2+^ and/or ± 5.6 mM glucose was added and incubated for 10 min or 1 hr at 16°C. Prior to imaging, the coverslip dishes were heated to 37 °C using a Heating Unit XL S (Zeiss) attached to the Zeiss AxioObserver Z1 (Carl Zeiss, Inc.). Images were collected under bright field illumination using a 40x C-Apochromat water immersion objective (N.A. 1.20), and ORCA-ER digital camera (Hamamatsu Photonics at 1 sec intervals for 5 min per field. The percentage of BAG1-mCherry positive bradyzoites displaying different types of gliding motility was calculate from 6 movies per sample. Images were imported into NIH ImageJ with a Cell Counter plug-in for quantification of the types of motility based on visual inspection.

### High-performance liquid chromatography UV (HPLC-UV) analysis of ATP, ADP and AMP levels in bradyzoites

BAG1-mCherry parasites were induced to form bradyzoites by culture at pH 8.2 in RPMI 1640 medium under ambient air (low CO_2_) for 7 days followed by scraping into ice-cold PBS containing 0.05% BSA. Cysts were released from host cells by repeated passage through a 23 *g* needle and collected by centrifugation at 150 *g* for 10 min. To purify bradyzoites, cysts were resuspended in 1 ml EC buffer without calcium or glucose but containing 10 μl biotinylated DBA (Vector laboratories) and 100 ul Pierce Streptavidin Magnetic Beads (Thermo Fisher) and incubated for 1 hr at 4°C. The beads and absorbed cysts were collected using a magnetic stand and resuspended in 1 ml EC buffer without calcium or glucose but containing 0.25 mg/ml trypsin and incubated for 10 min at 4°C. The supernatant containing released parasites was separated from the beads and retained. To remove any residual tachyzoites in the supernatant, 5 ul of mAb DG52 pre-coupled to 100 μl Dynabeads™ Protein G (Thermo Fisher) was added to the supernatant and incubated for 1 hr at 4°C. The supernatant was separated from the beads, bradyzoites centrifuged at 600 g, 4°C for 10 min, and resuspended in 1 ml EC buffer containing 1.8 mM Ca^2+^ and 5.6 mM glucose for 10 min or 1 hr at room temperature. Following incubation, the bradyzoites were pelleted at 600 g, 4°C for 10 mi and stored at −80°C until analysis.

A previously described method for extraction of ATP, ADP and AMP [83] was adapted for use here. In brief, 95 μl of extraction buffer (0.3 M perchloric acid (HClO_4_), 1 mM ethylenediaminetetraacetic acid disodium salt (Na_2_EDTA), pH 8.0) was used to resuspend cell pellets and incubated for 5 min at room temperature. Extraction was stopped by addition of 17 μl of neutralization buffer (2 M potassium hydroxide) to the samples followed by mixing. Samples were centrifuged at 14,000 *g* for 10 min at 4°C and the supernatant was transferred to a new tube for HPLC analysis. Analysis was performed using an HPLC system consisting of a SPD-20A UV/VIS detector (Shimadzu) equipped with SIL-20A autosampler (Shimadzu),with a Luna Omega Polar C18 column (4.6 mm internal diameter × 150 mm length, 3 μm particle size, 100 Å pore size), and LC-20AD pump (Shimadzu). The protocol was set up as isocratic separation using a mobile phase containing 0.1 M ammonium dihydrogen phosphate (NH_4_H_2_PO_4_, Sigma), pH 6.0, containing 1% methanol with a flow rate of 0.8 ml/min. Injection volume was 30 μl and peak detection was monitored at 254 nm. A series of standards containing ATP, ADP and AMP with different concentrations were used to establish retention times and standard calibration curves by calculating peak area. Samples from two independent biological replicates were analyzed using three technical replicates. The retention time and peak areas were used to calculate the corresponding concentration of each nucleotide from each sample according to the standard curve.

### Mouse infections and ex vivo cyst collection

Mice were housed in an Association for Assessment and Accreditation of Laboratory Animal Care International-approved facility at Washington University School of Medicine. All animal studies were conducted in accordance with the U.S. Public Health Service Policy on Humane Care and Use of Laboratory Animals, and protocols were approved by the Institutional Animal Care and Use Committee at the School of Medicine, Washington University in St. Louis.

Eight-week old female CD-1 mice (Charles River) were infected with 200 ME49 BAG1-mCherry GCaMP6f tachyzoites by intraperitoneal injection. After 30 days of infection, animals were sacrificed, the brain removed and homogenized and the number of brain cyst was determined by DBA staining and microscopy as previously described [77]. Eight-week old female CD-1 mice (Charles River) were infected with 5 cysts from the brain homogenate by oral gavage. Following a 30-day period these mice were euthanized, and brain homogenate was collected and added to glass bottom dishes for live imaging of tissue cysts.

### Statistical Analyses

Statistical analyses were performed in Prism (GraphPad). Data that passed normally distribution were analyzed by one-way ANOVA or Student’s t tests, while data that were not normally distributed, or contain too few samples to validate the distribution, were analyzed by Mann Whitney or Kruskal-Wallis non-parametric tests. *, *P* < 0.05, **, *P* < 0.01, ***, *P* < 0.001.

## Supporting information

Summplementary tables

## Acknowledgements

We thank Jennifer Powers Carson for technical help with the HPLC analysis which was performed in the Washington University Core Laboratory for Clinical Studies. We thank Vern Carruthers for providing plasmids, Louis Wiess and John Boothroyd for providing antibodies, members of the Sibley lab for helpful advice, Wandy Beatty, Microbiology Imaging Facility, for technical assistance with microscopy, and Jenn Barks for tissue culture support. Supported in part by a grant from the NIH (AI#034036).

## Author Contributions

Conceived and designed the experiments: Y.F., L.D.S.; Performed the experiments: Y.F.; Analyzed the data: Y.F., S.M., L.D.S.; Provided critical reagent and experimental advice: K.M.B., N.J., S.M.; Supervised the work S.M., L.D.S.; Wrote the manuscript: Y.F., L.D.S.; Edited the manuscript, all authors.

## Disclosures

The authors have no conflicts to disclose.

---

**Figure 4 figure supplement 1.**
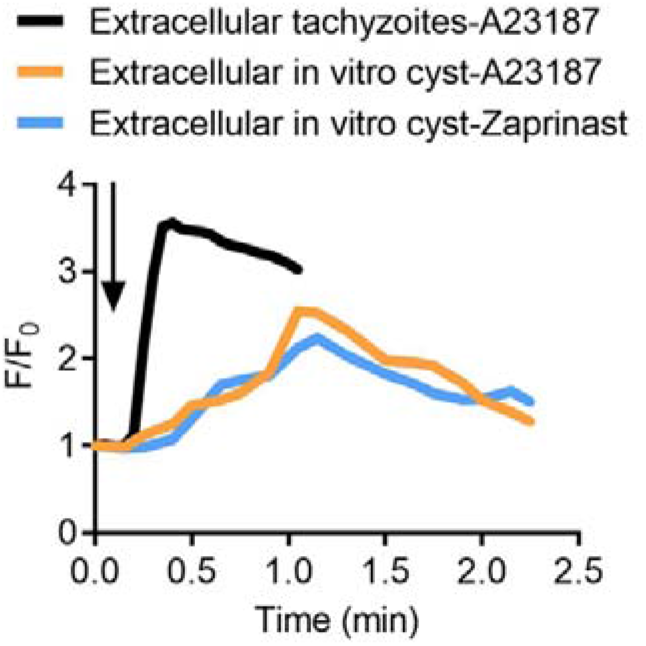
Calcium responses by extracellular tachyzoites and in vitro produced tissue cysts. **(A)** Fluorescence recording of ME49 strain parasites expressing GCaMP6f in response to A23187 (2 μM) or zaprinast (500 μM). Freshly harvested extracellular tachyzoites were compared to cysts induced in vitro in pH 8.2 RPMI 1640 medium for 7 days. Arrow indicates time of addition of calcium agonists. Each kinetic curve represented the mean of 3 independent samples.

**Figure 6 figure supplement 1.**
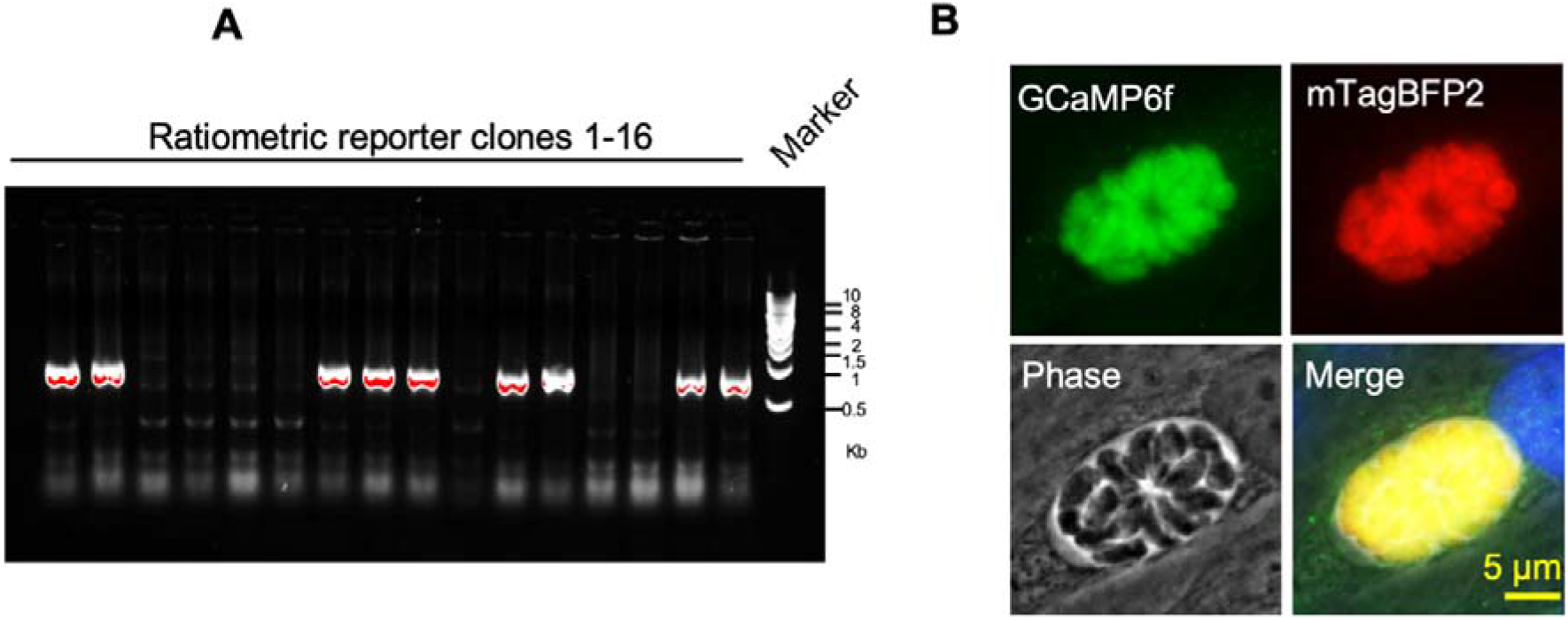
figure supplement 1 Identification of ME49 GCaMP6f-P2A-mTagBFP2 BAG1-mCherry ratiometric reporter by PCR and IFA. **(A)** Transgenic screening of clones of ME49 GCaMP6f BAG1-mCherry parasites expressing P2A-mTagBFP2 at the C-terminal of GCaMP6f using PCR amplification with primer set P1-P2 shown in diagram in **Figure 6A**. **(B)** IFA analysis showing co-localization of GCaMP6f and mTagBFP2 in tachyzoites of the dual reporter strain grown in HFF cells for 24 hr. Monoclonal anti-His antibody was used to stain GCaMP6f while rabbit anti-tRFP antibody was used to stain mTagBFP2 followed by goat anti-mouse IgG conjugated to Alexa Fluor-488 and goat anti-rabbit IgG conjugated to Alexa Fluor-568 secondary antibodies. Scale bar = 5 μm.

**Figure 8 figure supplement 1.**
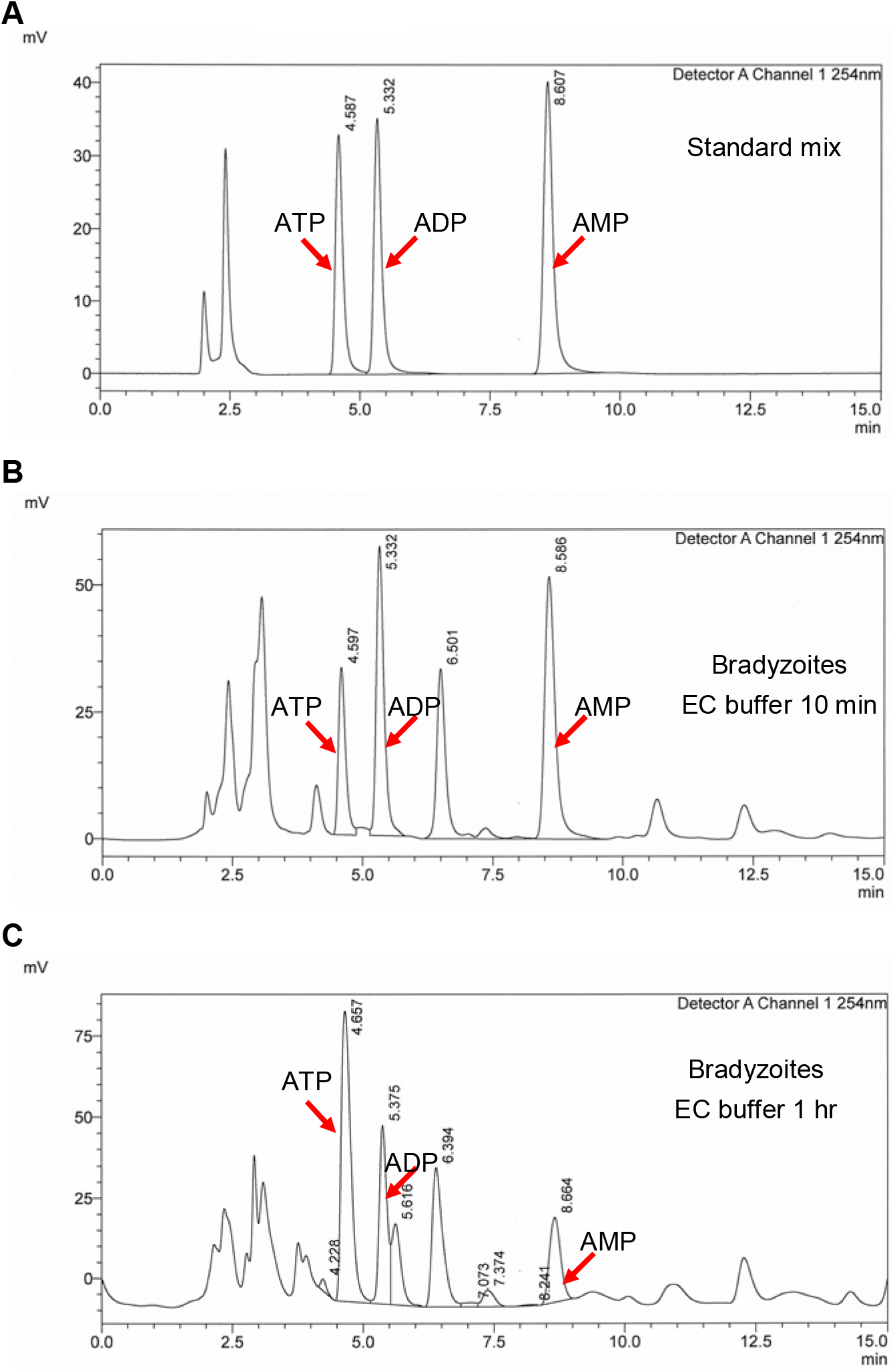
figure supplement 1 Establishment of HPLC-UV analysis of ATP, ADP and AMP levels in parasites. **(A)** HPLC-UV chromatograms of standard mix containing 12.5 μM ATP, 12.5 μM ADP and 12.5 μM AMP. Arrows indicate the peaks of ATP, ADP and AMP. **(B)** HPLC-UV chromatograms of ATP, ADP and AMP extracts from bradyzoites (2 x10^7^) incubated with EC buffer containing 1.8 mM Ca^2+^ and 5.6 mM glucose for 10 min. Arrows indicate the peaks of ATP, ADP and AMP. **(C)** HPLC-UV chromatograms of ATP, ADP and AMP extracts from bradyzoites (1.2 × 10^7^) incubated with EC buffer containing 1.8 mM Ca^2+^ and 5.6 mM glucose for 1 hr. Arrows indicate the peaks of ATP, ADP and AMP.

## Rich Media Files

**Figure 1-video 1 Egress by ME49 BAG1-mCherry tachyzoites in response to A23187.**

Time-lapse video microscopy showing A23187 (2 μM) induced egress of ME49 BAG1-mCherry strain tachyzoites grown in vitro in HFF cells for 24 hr. Videos for intracellular tachyzoites in EC buffer were recorded for 10 min and A23187 (2 μM) was added 30 s after the recording initiated. Display frame rate is 8 frames per second while the acquisition frame rate is 3 frames per second. Bar = 10 μm.

**Figure 1-video 2 Egress by ME49 BAG1-mCherry bradyzoites in response to A23187.**

Time-lapse video microscopy showing A23187 (2 μM) induced egress of ME49 BAG1-mCherry strain bradyzoites induced by in vitro culture on HFF cells for 7 days at pH 8.2. Videos for intracellular bradyzoites in EC buffer were recorded for 10 min and A23187 (2 μM) was added 30 s after the recording initiated. Display frame rate is 4 frames per second while the acquisition frame rate is 10 frames per second. Bar = 10 μm.

**Figure 2-video 1 A23187 - induced permeabilization of the parasitophorous vacuole membrane (PVM) detected by vacuolar leakage of FNR-mCherry secreted by tachyzoites.**

Time-lapse video microscopy showing A23187 (2 μM)-induced FNR-mCherry leakage from the PV surrounding FNR-mCherry BAG1-EGFP expressing tachyzoites. FNR-mCherry BAG1-EGFP tachyzoites cultured under normal condition in HFF cells for 24 hr were treated with A23187 (2 μM) in EC buffer for 10 min at 37℃. Videos were recorded for 10 min and A23187 (2 μM) was added 30 s after the recording initiated. Display frame rate is 6 frames per second while the acquisition frame rate is 5 frames per second. Bar = 5 μm.

**Figure 2-video 2 Trypsin - induced disruption of in vitro differentiated tissue cysts expressing ME49 FNR-mCherry BAG1-EGFP.**

Time-lapse video microscopy showing A23187-induced FNR-mCherry leakage in vitro differentiated tissue cysts of FNR-mCherry BAG1-EGFP bradyzoites. FNR-mCherry BAG1-EGFP bradyzoites induced by cultivation in HFF cells in vitro for 7 days at pH 8.2 were treated with 0.25 mg/ml Trypsin in EC buffer for 6 min at 37°C. Videos were recorded for 6 min and 0.25 mg/ml Trypsin was added 30 s after the recording initiated. Display frame rate is 3 frames per second while the acquisition frame rate is 15 frames per second. Bar = 5 μm.

**Figure 2-video 3 A23187 -induced permeabilization of in vitro differentiated tissue cysts detected by vacuolar FNR-mCherry leakage from ME49 FNR-mCherry BAG1-EGFP bradyzoites.**

Time-lapse video microscopy showing A23187 (2 μM)-induced FNR-mCherry leakage from in vitro differentiated cysts of FNR-mCherry BAG1-EGFP. FNR-mCherry BAG1-EGFP bradyzoites induced by cultivation in HFF cells in vitro for 7 days at pH 8.2 were treated with A23187 (2 μM) in EC buffer for 10 min at 37°C. Videos were recorded for 10 min and A23187 (2 μM) was added 30 s after the recording initiated. Display frame rate is 3 frames per second while the acquisition frame rate is 15 frames per second. Bar = 5 μm.

**Figure 3-video 1 Calcium response of ME49 BAG1-Cherry GCaMP6f expressing tachyzoites stimulated by A23187.**

Time-lapse video microscopy showing GCaMP6f fluorescence changes of intracellular ME49 BAG1-mCherry GCaMP6f tachyzoites grown in HFF cells in vitro for 24 hr in response to A23187 (2 μM) in EC buffer. Videos were recorded for 10 min and A23187 (2 μM) was added 30 s after the recording initiated. Display frame rate is 10 frames per second while the acquisition frame rate is 3 frames per second. Bar = 10 μm.

**Figure 3-video 2 Calcium response of ME49 BAG1-Cherry GCaMP6f expressing bradyzoites stimulated by A23187.**

Time-lapse video microscopy showing GCaMP6f fluorescence changes of intracellular ME49 BAG1-mCherry GCaMP6f bradyzoites induced by cultivation in HFF cells in vitro for 7 days at pH 8.2 in response to A23187 (2 μM) in EC buffer. Videos were recorded for 14 min and A23187 (2 μM) was added 30 s after the recording initiated. Display frame rate is 6 frames per second while the acquisition frame rate is 10 frames per second. Bar = 10 μm.

**Figure 4-video 1 Calcium response of ME49 BAG1-mCherry GCaMP6f cysts isolated from chronically infected mouse brains and treated in vitro with DMSO.**

Time-lapse video microscopy showing GCaMP6f fluorescence changes of ME49 BAG1-mCherry GCaMP6f cysts isolated 30 days post-infection from the brains of chronically infected mice in response to DMSO (0.1%) in EC buffer. Videos were recorded for 5 min and DMSO (0.1%) was added 15 s after the recording initiated. Display frame rate is 6 frames per second while the acquisition frame rate is 3 frames per second. Bar = 2 μm.

**Figure 4-video 2 Calcium response of ME49 BAG1-mCherry GCaMP6f cysts isolated from chronically infected mouse brains and treated in vitro with A23187.**

Time-lapse video microscopy showing GCaMP6f fluorescence changes of ME49 BAG1-mCherry GCaMP6f cysts isolated 30 days post-infection from chronically infected mice in response to A23187 (2 μM) in EC buffer. Videos were recorded for 5 min and A23187 (2 μM) was added 15 s after the recording initiated. Display frame rate is 6 frames per second while the acquisition frame rate is 5 frames per second. Bar = 2 μm.

**Figure 5-video 1 Calcium response of extracellular ME49 BAG1-mCherry GCaMP6f tachyzoite in response to A23187.**

Time-lapse video microscopy showing GCaMP6f fluorescence changes of extracellular ME49 BAG1-mCherry GCaMP6f tachyzoite in response to A23187 (2 μM) in EC buffer. Videos were recorded for 3 min and A23187 (2 μM) was added 15 s after the recording initiated. Display frame rate is 4 frames per second while the acquisition frame rate is 3 frames per second. Bar = 2 μm.

**Figure 5-video 2 Calcium response of extracellular ME49 BAG1-mCherry GCaMP6f bradyzoite in response to A23187.**

Time-lapse video microscopy showing GCaMP6f fluorescence changes of extracellular ME49 BAG1-mCherry GCaMP6f bradyzoite in response to A23187 (2 μM) in EC buffer. Bradyzoites were liberated by 0.25 mg/ml trypsin for 5 min from in vitro cysts induced for cultivation in HFF cells for 7 days at pH 8.2. Videos were recorded for 3 min and A23187 (2 μM) was added 15 s after the recording initiated. Display frame rate is 2 frames per second while the acquisition frame rate is 5 frames per second. Bar = 2 μm.

**Figure 7-video 1 Trypsin induced liberation of ME49 BAG1-mCherry GCaMP6f bradyzoites from in vitro cultured cysts.**

Time-lapse video microscopy recording GCaMP6f fluorescence changes from BAG1-mCherry GCaMP6f bradyzoites induced by cultivation in HFF cells for 7 days at pH 8.2 during digestion by 0.25 mg/ml Trypsin in EC buffer. Videos were recorded for 6 min and 0.25 mg/ml trypsin was added 30 s after the recording initiated. Display frame rate is 16 frames per second while the acquisition frame rate is 5 frames per second. Bar = 5 μm.

**Figure 7-video 2 Gliding motility of ME49 BAG1-mCherry GCaMP6f bradyzoites released from in vitro cysts.**

Time-lapse video microscopy of gliding motility of bradyzoites liberated by 0.25 mg/ml trypsin for 5 min from in vitro cyst induced by cultivation in HFF cells for 7 days at pH 8.2. Images were collected using spinning disc confocal microscopy. The arrow shows the gliding motility of bradyzoite in EC buffer. Videos were recorded for 2 min. Display frame rate is 6 frames per second while the acquisition frame rate is 1 frame per second. Bar = 2 μm.

## Supplementary Files

Supplementary Table 1: Primers used in this study

Supplementary Table S2 Plasmids used in this study

Supplementary Table S3 Parasite lines used in this study

## Notes

### Competing Interest Statement

The authors have declared no competing interest.

### Summary of Updates

We have added to data in Figure 8 to address the energy levels of bradyzoites and their ability to respond to changes in the environment.

