## Supplementary material for "Toxoplasma bradyzoites exhibit physiological plasticity of calcium and energy stores controlling motility and egress": Summplementary tables: Supplementary table 2 Parasite lines and plasmids.docx

Supplementary Table S2 Plasmids used in this study

| Plasmid | Usage | Source |
| --- | --- | --- |
| pBAG1:mCherry, SAG1:CAT, TUB1:GCaM6f (pNJ-26) | Generation of BAG1-mCherry GCaMP6f line | This study (Nathaniel Jones) |
| pSAG1:CAS9-EGFP, U6:sgUPRT | Template for construction of pSAG1:CAS9-GFP, U6:sgDHFR 3’UTR | Shen et al., 2014 (1) |
| pSAG1:CAS9-EGFP, U6:sgDHFR 3’UTR | Generation of ratiometric reporter | This study |
| p2A-mTagBFP2, DHFR-TS:HXGPRT | Generation of ratiometric reporter | This study |
| pBAG1:EGFP, DHFFR-TS::HXGPRT | Generation of BAG1-EGFP line | This study |
| pBAG1:mCherry, DHFFR-TS::HXGPRT | Generation of BAG1-mCherry line | This study |
| pMIC2:GLuc-myc, DHFR-TS | Generation of MIC2 secretion reporter BAG1-mCherry MIC2-GLuc | Brown et al., 2016 (2) |
| pTUB1:FNR-mCherry, CAT | Generation of BAG1-EGFP FNR-mCherry line | Carruthers lab |
| pTUB1:YFP-mAID-3HA, DHFR-TS:HXGPRT | Template for construction of p2A-mTagBFP2, DHFR-TS:HXGPRT, pBAG1:EGFP, DHFFR-TS::HXGPRT and pBAG1:mCherry, DHFFR-TS::HXGPRT | Brown et al., 2017 (3) |

1.	Shen B, Brown K, Long S, & Sibley LD (2017) Development of CRISPR/Cas9 for Efficient Genome Editing in *Toxoplasma gondii*. *Methods in molecular biology* 1498:79-103.

2.	Brown KM, Lourido S, & Sibley LD (2016) Serum Albumin Stimulates Protein Kinase G-dependent Microneme Secretion in Toxoplasma gondii. *The Journal of biological chemistry* 291(18):9554-9565.

3.	Brown KM, Long S, & Sibley LD (2017) Plasma Membrane Association by N-Acylation Governs PKG Function in Toxoplasma gondii. *mBio* 8(3).
