## Supplementary material for "Toxoplasma bradyzoites exhibit physiological plasticity of calcium and energy stores controlling motility and egress": Summplementary tables: Supplementary Table S3 Parasite lines used in this study.docx

| Line | Genotype | Source |
| --- | --- | --- |
| ME49 Δ*hxgprt::FLUC* | ME49 Δ*hxgprt::TUB1:FLUC* | Tobin et al., 2012 (Tobin and Knoll 2012) |
| BAG1-mCherry GCaMP6f | ME49 Δ*hxgprt*::*TUB1:FLUC*; *BAG1*:*mCherry*, *SAG1*:*CAT*, *TUB1*:*GCaMP6f* | This study |
| BAG1-mCherry | ME49 Δ*hxgprt*::*TUB1:FLUC*; *BAG1*:*mCherry*, *DHFR-TS*:*HXGPRT* | This study |
| BAG1-EGFP | ME49 Δ*hxgprt*::*TUB1:FLUC*; *BAG1*:*EGFP*, *DHFR-TS*:*HXGPRT* | This study |
| BAG1-mCherry MIC2-GLuc | ME49 Δ*hxgprt*::*TUB1:FLUC*; *BAG1*:*mCherry*, *DHFR-TS*:*HXGPRT; MIC2*:*MIC2-GLuc, DHFR-TS* | This study |
| BAG1-EGFP FNR-mCherry | ME49 Δ*hxgprt*::*TUB1:FLUC*; *BAG1*:*EGFP*, *DHFR-TS*:*HXGPRT ; SAG1*:*CAT*, *TUB1:FNR-mCherry* | This study |
| BAG1-mCherry GCaMP6f-P2A-mTagBFP2  (Ratiometric reporter) | ME49 Δ*hxgprt*::*TUB1:FLUC*; *BAG1*:*mCherry*, *SAG1*:*CAT*, *TUB1*:*GCaMP6f-P2A-mTagBFP2, DHFR-TS*:*HXGPRT* | This study |

Tobin, C. M. and L. J. Knoll (2012). "A patatin-like protein protects Toxoplasma gondii from degradation in a nitric oxide-dependent manner." Infect Immun **80**(1): 55-61.
